# DExCon, DExogron, LUXon: on-demand expression control of endogenous genes reveals differential dynamics of Rab11 family members

**DOI:** 10.1101/2021.12.03.471086

**Authors:** Jakub Gemperle, Thomas Harrison, Chloe Flett, Antony Adamson, Patrick Caswell

## Abstract

CRISPR technology has made generation of gene knockouts widely achievable in cells. However, once inactivated, their reactivation remains difficult, especially in diploid cells. Here, we present DExCon (Doxycycline-mediated endogenous gene Expression Control), DExogron (DExCon combined with auxin-mediated targeted protein degradation) and LUXon (light responsive DExCon), approaches which combine one-step CRISPR-Cas9 mediated targeted knock-in of fluorescent proteins with an advanced Tet-inducible TRE3GS promoter. These approaches combine blockade of active gene transcription with the ability to reactivate transcription on demand, including activation of silenced genes. Systematic control can be exerted using doxycycline or spatiotemporally by light, and we demonstrate functional knockout/rescue in the closely related Rab11 family of vesicle trafficking regulators. Fluorescent protein knock-in results in bright signals compatible with low-light live microscopy from monoallelic modification, the potential to simultaneously image different alleles of the same gene and bypasses the need to work with clones. Protein levels are easily tunable to correspond with endogenous expression through cell sorting (DExCon), timing of light illumination (LUXon) or by exposing cells to different levels of auxin (DExogron). Furthermore, our approach allowed us to quantify previously unforeseen differences in vesicle dynamics, expression kinetics and protein stability among highly similar endogenous Rab11 family members and their colocalization in triple knock-in cells.

**GRAPHICAL ABSTRACT:** 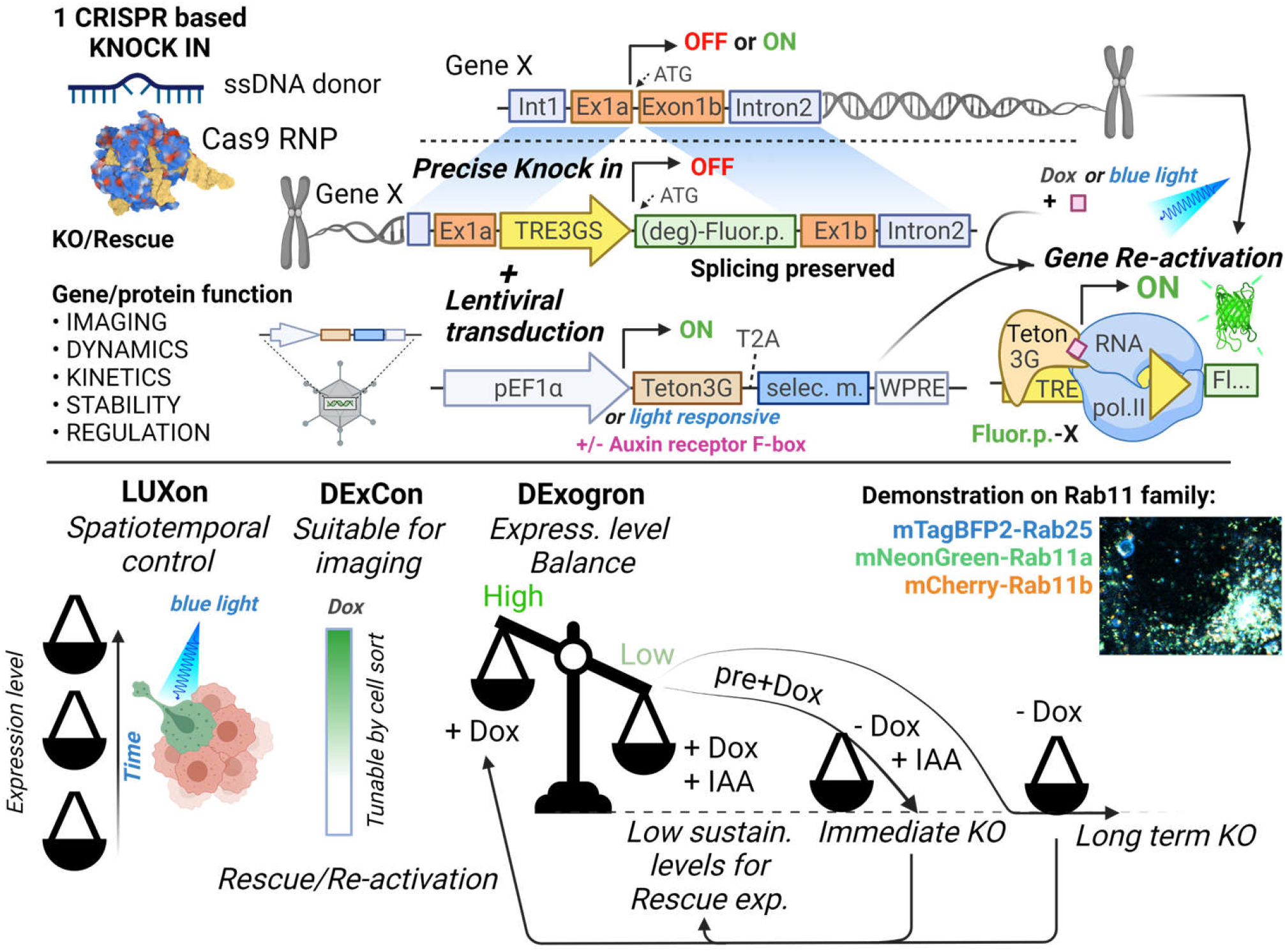

**IN BRIEF:** We describe development of DExCon, LUXon and DExogron approaches, where a single CRIPR/Cas9-mediated gene editing event can block endogenous gene expression, with the ability to reactivate expression encoded such that even silent genes can be expressed. Expression can be controlled systematically using doxycycline, or spatiotemporally by light, allowing fluorescent tagging of endogenous proteins and quantification of expression kinetics, protein dynamics and stability for highly similar genes such as members of the Rab11 family.

## Introduction

The ability to control gene expression and analyse protein coding gene function inside cells is a critical step in our understanding of normal physiology and disease pathology. CRISPR/Cas9 technology has revolutionised gene editing approaches, and makes generation of gene knockouts or knock-ins straightforward^1^. Combination of CRISPR with single-stranded DNA (ssDNA) donors has proven to be the most precise and efficient method of on-target integration^2,3^, although knock in approaches still suffer from low efficiencies, especially when aiming to achieve homozygosity^4^. Fluorescent protein (FP) knock in at endogenous loci leading to protein-fusion allows functional enrichment of complete knock-in events and study of native protein function using fluorescent microscopy^5,6^. However, they only complement loss-of-function/gain-of-function methods and their usefulness is limited by brightness of the chosen FP. Conditional loss-of-function methods mainly rely on CRISPR/Cas9 combination with FLP-frt or Cre-LoxP systems^2^. However, these are labour-intensive and mediated by recombination, which is by nature irreversible thus complicating attempts to rescue gene expression. In addition, to understand the full spectrum of gene functions, it is advantageous to reversibly tune expression levels, visualized by fluorescence microscopy, in a spatio-temporal context and not only depend on binary on/off approaches. Such levels of control can be partly offered by the auxin-inducible degron (AID) system^7^ introduced as a tag into the endogenous locus via CRISPR/Cas9^8^. Here, an exogenously expressed plant auxin receptor F-box protein triggers ubiquitylation and degradation of target proteins fused to an AID tag on addition of the plant hormone auxin. However, this technology has its own limitations and caveats; for example, poor spatial control due to diffusion of auxin, significant basal degradation without auxin and/or incomplete auxin-induced protein depletion.

Ideally, gene expression control and visualization of its protein product should be generated by a single gene modification (e.g. CRISPR knock in) at the endogenous locus and allow temporal, conditional and reversible gene inactivation of all protein-coding transcripts, on demand, without the drawbacks of AID system. Such an approach is particularly useful for studying highly similar protein coding genes where their complementary and specific functions remain poorly understood, such as for members of Rab11 family of small GTPases: Rab11a, Rab11b, and Rab25 (Rab11c)). Rab11s share high amino acid identity (Rab11a:Rab11b 89 %; Rab11a:Rab25 66%; Rab11b:Rab25 61%), are known to play key roles in membrane transport, localize to recycling endosomes and have been identified as important players in the cellular basis of an ever-increasing number of human disorders, including cell migration/invasion and cancer^9,10^. Several Rab11 family isoforms have been reported^11^ and specific antibodies are hard to obtain without crossreactivity. Moreover, much what is known about their regulation and dynamics in live cells has been achieved with the aid of transiently transfected cDNA-mediated overexpression which may lead to non-physiological artefacts^12,13^.

To overcome limitations of previously established gene expression control technologies, we generated tractable single CRISPR knock-in based strategies for tunable and reversible gene expression control capable of knock-out/rescue and endogenous gene re-activation. These modifications are ideal for microscopy when combined with FP knock-in, are free of transfection artefacts, preserve posttranscriptional and posttranslational control and avoid comprehensive and labour-intensive genotyping post-editing of clones. Specifically, we harnessed CRISPR/Cas9 technology for targeted knock-in of the third generation Tetracycline-Inducible System (Tet-On 3G) promoter TRE3GS followed by a FP coding region. This promoter lacks binding sites for endogenous mammalian transcription factors, is effectively silent in the absence of induction and any transcription driven by endogenous active promoter generates nonsense frameshift. Simultaneous or sequential silencing of additional alleles was achieved by additional ssDNA donor coding antibiotic resistance to attain complete and selectable functional knockout. Spatiotemporal control was achieved by combination with a photoactivatable (PA)-Tet-OFF/ON system^14^, even in cells within 3D microenvironments. We further established a tunable dual-input system combining TetON and AID, termed DExogron, that enhanced switch off kinetics of the Tet ON system, allowing rapid complete protein depletion or adjustment of expression to physiological levels on demand.

Our optimised pipeline to deliver this suite of CRISPR tools was applied to modulate endogenous gene expression of Rab11 family members, demonstrating that re-activation of endogenous Rab25 expression can promote the invasive migration of cancer cells, and revealing new insight into the localisation and functions of Rab11a, Rab11b and Rab25.

## Results

Deconvolving the relative functions of genes within highly related gene families has proven difficult, for example the Rab11 family has a complex relationship with cancer, with several potential oncogenic and tumour suppressive functions identified^10,15,16^. However, gene editing offers potential solutions. In order to identify the best knock in strategies to visualize the Rab11 gene family, we first analysed expression patterns of the Rab11s across tissues and in cancer using UCSC Xena, an online exploration tool for multi-omic data^17^. Rab11a, Rab11b and Rab25 were significantly enriched in ovarian cancer, both for primary and recurrent tumours (S1A). Bioinformatic analyses suggested the existence of additional potentially protein-coding human Rab11a/b splice variants (S1B). Analysis of expression quantitative trait loci (eQTLs) revealed abundant expression of two Rab11b- and five Rab11a-coding transcripts across multiple healthy tissues while Rab25 is more restricted, but expression is increased across multiple types of cancer (S1C, https://doi.org/10.48420/16988617). Similarly, Rab11a/Rab11b /Rab25 splicing isoforms are enriched in ovarian cancer compared to normal tissue (S1D). All significantly expressed transcripts share an intact N-terminus, however for some splice variants, amino acids important for GTP/GDP binding (which allow oscillation between active/inactive states) or for the attachment to the outer leaflet of endosomal membranes are missing, which could therefore profoundly alter Rab11s function (S1E). Thus, to capture the full physiological picture of the combination of Rab11s protein coding transcripts, we fluorescently tagged the N-terminus of Rab11s using CRISPR knock in (1A). We used long single-stranded DNA (ssDNA) donor for precise and efficient on-target integration^2,3,18^ with flanking homologous arms 150-300 bases prepared via RNA intermediates (1B). Initial testing of different approaches led to our optimized protocol (1C) combining magnetofection of long ssDNA and high fidelity (Hifi) Cas9 delivered as ribonucleoprotein complex with guideRNA (crRNA:tracrRNA). This combination reduces off target effects to non-detectable levels and leads to high-fidelity hetero/homozygous knock-in^19^. To avoid comprehensive and labour-intensive genotyping post-editing of clones^18^, we took the advantage of fluorescent readouts to monitor and enrich correct homology directed repair (HDR) by FACS sorting^3,6^. Pilot testing led to a high rate of successful knock in HEK293T cells with expected localisation of Rab11a (10-20 %) or Rab11b (2-10 %), respectively (S1F). For further work we selected a commonly used cell line model for ovarian cancer, A2780, and knocked in mNeonGreen or mCherry after the start codon of Rab11a or Rab11b, respectively (1D). This integration, although less efficient (0.2-1 %) compared to HEK293T, was specific as all sorted cells exhibited correct Rab11a/b localization and our protocol was also successful in a second ovarian cancer cell line that has defect in homologous recombination (BRCA-1 mutant COV362^20^; 1E). Western blotting with antibodies that recognise Rab11a, or both Rab11a and Rab11b, confirmed knock-in (1F). Consistent with the literature^3,21^, a proportion of cells showed truncation of the linker flanking the FP (1G), but not in the integrated homologous arms, indicating that sorting cells for fluorescence (or another functional protein) is important for establishing in frame knock-in at the N-terminus preserving protein function. This potential caveat was demonstrated by our effort to knock in an additional non-fluorescent component, antiGFPnanobody, together with mCherry. Although knock in to the Rab11a locus showed the expected localization of mCherry-Rab11a in all sorted cells (S2A), integration of the antiGFPnanobody sequence was not complete in ~90% of mCherry-positive cells (S2B-C).

**Figure 1.**
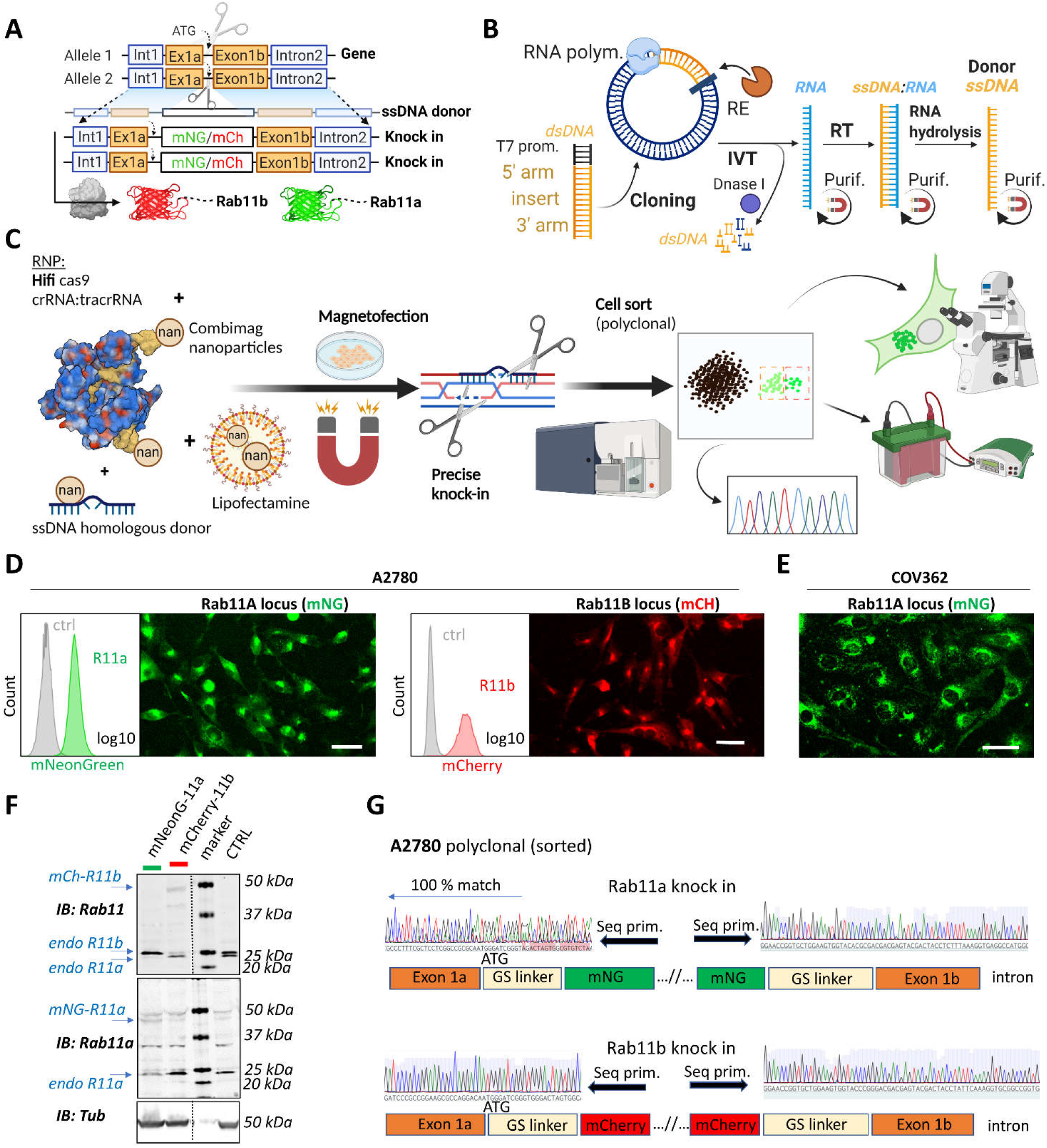
A rapid and efficient pipeline for CRIPR/Cas9 knock-in of fluorescent proteins at the endogenous locus of Rab11 family members. A-C) Schematic diagram of ssDNA mediated knock in strategy of different fluorophores with corresponding homologous arms A), the ssDNA preparation technique (IVT = in vitro transcription; RT = reverse transcription; RE = restriction endonuclease; purif. = purification using magnetic SPRI beads) B) and our optimized pipeline (RNP = ribonucleoprotein complex) C). Illustrations were created with BioRender.com. FACS and fluorescence widefield images (background subtracted) of mNeonGreen or mCherry knock ins to Rab11A or Rab11b loci of D) A2780 or E) COV362 ovarian cancer cell lines. Ctrl represent un-modified cells; scale bar=40 μm. F) Immunoblots of mNeonGreen-Rab11a, mCherry-Rab11b and un-modified (Ctrl) cells. Fluorescent antibodies, specific anti-Rab11a or antibody targeting both Rab11a/b (Rab11) shown as black and white. Tubulin, loading control. G) Chromatograms of mNeonGreen-Rab11a and mCherry-Rab11b cells with schematic of knock in outcome (seq. prim = sequencing primer).

### DExCon supresses endogenous gene expression and re-activates in a dox-dependent manner

Being able to visualize endogenous Rab11a/b, we analysed sequences of different promoters for their ability manipulate gene expression at the endogenous locus. The most promising candidate was the advanced non-leaky TRE3GS promoter of the third generation Tetracycline-inducible System (Tet-On 3G) as it offered conditional and reversible control regulated by the small molecule doxycycline (dox). Analysis of 365 bp TRE3GS promoter revealed four ATG codons, all of which could be designed out of frame to the endogenous start codon, leading to multiple premature stop codons in transcripts driven by basal expression of the endogenous active promoter (2A). Tet-On 3G transactivator was stably delivered to A2780 cells by lentivirus in tandem with mTagBFP (separated by the selfcleaving peptide T2A) and cells selected for mTagBFP fluorescence. The TRE3GS promoter sequence was introduced into the donor sequence upstream of mNeonGreen or mCherry and delivered by long ssDNA mediated knock in to the Rab11a or Rab11b locus, respectively, in A2780 cells. Sorted cells showed expression of corresponding FP-Rab11 with a perinuclear vesicle-enriched localisation matching the endogenous protein, but only in the presence of dox, and decreased levels of non-modified endogenous Rab11a/b indicating that DExCon blocks expression at the endogenous locus (2B, C; S3A).

**Figure 2.**
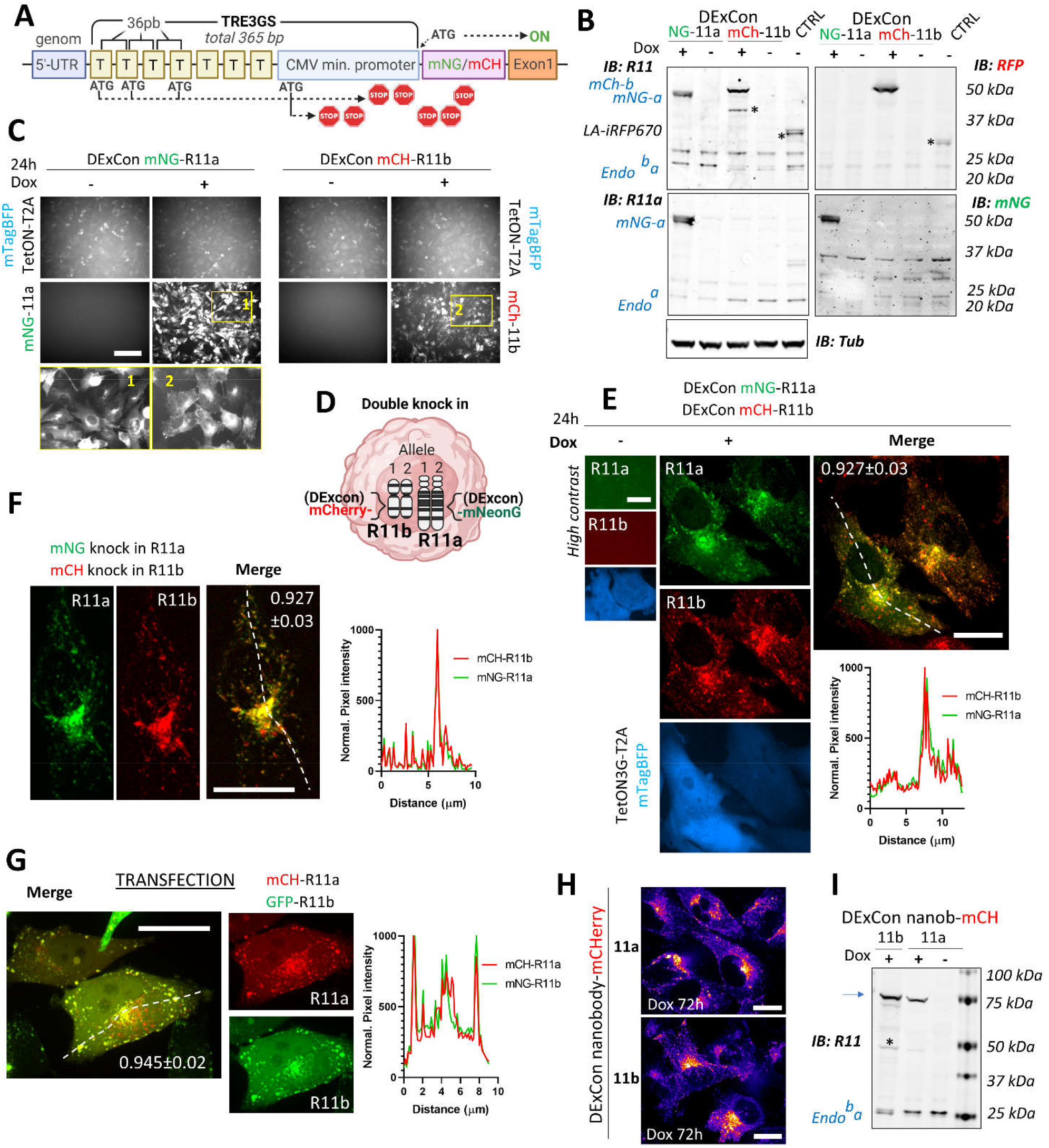
Reversible suppression of endogenous gene expression with DExCon. A) DExCon schematic. B) DExCon knock in cells were dox treated for 24-72h, sorted for mCherry or mNeonGreen fluorescence and grown for 2 weeks without dox before re-analysis. Representative immunoblots of mNeonGreen-Rab11a or mCherry-Rab11b DExCon A2780 cells treated or not treated with dox for 48h (a = Rab11a; b = Rab11b) probed with anti-mNeonGreen (mNG), anti-mCherry (RFP), anti-Rab11a or anti-Rab11 (targeting both Rab11a/b) antibodies shown as black and white. Tubulin (Tub), loading control. CTRL represents wt A2780 over-expressing Lifeact-iRFP670. Stars indicate mCherry/iRFP670 lower molecular weight band caused by hydrolysis during sample preparation^22^. For higher contrast see S3A. C) Live fluorescence images of mNeonGreen-Rab11a or mCherry-Rab11b DExCon (A2780) cells ± dox treatment (24h). Scale bar=100 μm. Area in yellow rectangle (1 or 2) is shown at the bottom with higher resolution. D) Schematic of double knock in, classical or DExCon, within same cell. E-G) Spinning disk confocal images of E) double DExCon Rab11a/b expressing Teton3G-T2A-mtagBFP ± dox, F) classical double mNeonGreen or mCherry knock ins to Rab11a or Rab11b as indicated and G) A2780 co-transfected (magnetofection) with cDNA for GFP-Rab11b and mCherry-Rab11a. Maximum intensity z-projection are shown, scale bar=20μm. Plot profiles correspond to the dashed line and numbers reflect Pearson cross correlation coefficient (average ± SEM, n = 14-22 cells). Spinning disk confocal images H) or immunoblots I) of antiGFPnanobody-mCherry DExCon cells (mCherry channel as “gem” LUT), modified in Rab11a or Rab11b locus treated ± dox for 72h. Scale bar=20μm. Blue arrow points to the full-length fusion product.

To exemplify the versatility of our CRISPR knock-in pipeline, we generated double mNeonGreen / mCherry knock-in at Rab11a /Rab11b loci respectively with and without the DExCon module (2E, F). In all combinations, endogenous Rab11a/Rab11b localisation driven by dox-induced or endogenous promoter were significantly correlated, and both Rab11a and Rab11b concentrated in vesicles at the perinuclear recycling compartment (2E, F). By contrast, transient transfection of plasmids coding FP-Rab11a/Rab11b gave very bright overlapping signals, but large vesicles were observed particularly towards the cell periphery (2G). These findings were confirmed when red/green-FPs were switched (S3B and S3C) and suggest that DExCon can induce expression from the endogenous gene locus that preserves physiological distribution of the protein within cells with signal intensities compatible with low light/exposure live microscopy (S3D; movie S1-2). This was further demonstrated by imaging Rab11a/b vesicles transporting the recognised cargo transferrin (S3E; movie S3) and by imaging cells in 3D environments. In 3D-cell derived matrix (CDM), higher exposures were required for visualisation of conventional double knock-ins, and this led to a lag in capture between channels and significant bleaching of mCherry over time (S3F; movie S4). However, DExCon knock-ins showed higher intensity and could therefore be visualised at lower exposure, avoiding caveats associated with lag (mislocalisation) and photobleaching, revealing that whilst Rab11a vesicle intensity is consistent within polarised cells, Rab11b intensity is increased at the tips of protrusions relative to the region of the cell in front of the nucleus (S3G).

We next explored the ability of DExCon to deliver full length knock in of GFPnanobody-mCherry sequence to Rab11 gene loci as its flanking by an upstream TRE3GS sequence could improve the likelihood of complete integration and positive selection. Upon dox induction, all sorted cells exhibited the expected localization of Rab11a or Rab11b (2H) and fulllength size on western blot (2I) indicating that encoding functionality at each of the 5’ and 3’ ends of ssDNA donor allows enrichment of accurately modified cells even for non-fluorescent coding genes.

### Reactivation of silenced genes by DExCon and LUXon

A2780 ovarian cancer cells show abundant Rab11a/b expression while the more restricted family member Rab25 is not expressed (3A). To study endogenous Rab25 in this cell line, we hypothesised that our DExCon strategy could reactivate Rab25 expression from the genome (3B). TRE3GS-mNeonGreen was delivered by long ssDNA mediated knock-in to the Rab25 locus in A2780 cells expressing Tet-On 3G transactivator. Dox treatment led to mNeonGreen fluorescence with expected perinuclear localization of mNeonGreen-Rab25 in all sorted cells (in 2D and 3D-CDM; 3C) and specificity of integration was confirmed by western blot (S4A) and sequencing (S4B).

**Figure 3.**
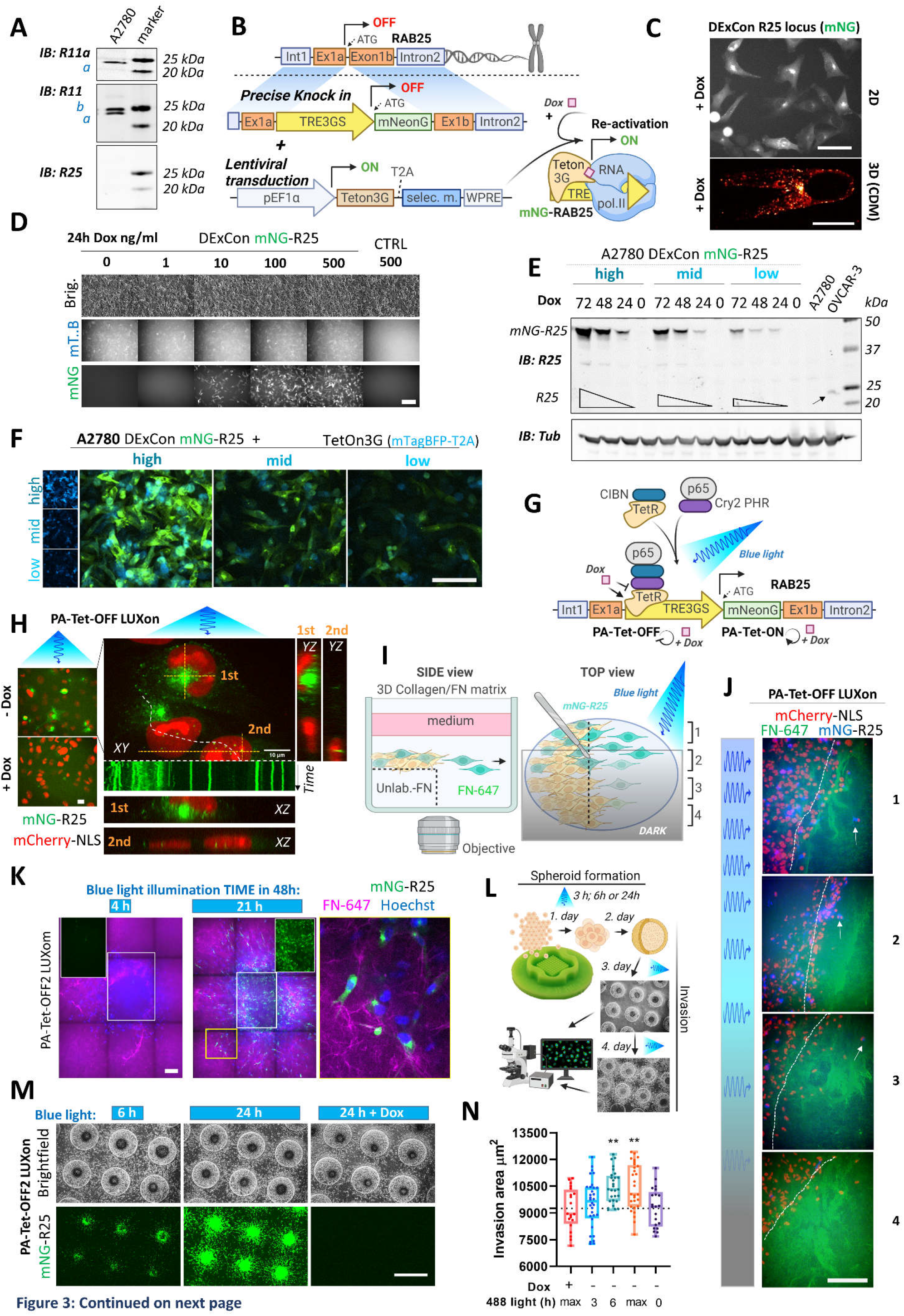
Spatiotemporal control of gene re-activation with DExCon and LUXon. A) Immunoblots of A2780 cells lysates probed with antibodies specific for anti-Rab11a, targeting both Rab11a/b (Rab11) or Rab25 (a = Rab11a; b = Rab11b). Tubulin, loading control. B) Schematic of the DExCon mNeonGreen knock in strategy and lentiviral transduction for dox dependent re-activation of Rab25 expression. C-D) Rab25 DExCon knock in cells were dox treated for 24-72h, sorted for mNeonGreen fluorescence followed by cell growth 2 weeks without dox before re-analysis. Live fluorescence images of re-activated Rab25 fused to mNeonGreen (mNG) in A2780 24h after dox treatment (250 ng/ml). C) Top: Cells on tissue culture-treated plastic (scale bar=100 μm). Bottom: Cell in CDM (Spinning disc confocal image; scale bar=20μm). D) Cells exposed to increasing dox concentration imaged by brightfield and fluorescence microscopy (mT..B = Teton3G-T2A-mTagBFP; brig.=brightfield; Ctrl = un-modified cells). Scale bar=100 μm. E) Immunoblots of mNeonGreen-Rab25 DExCon re-sorted cells as indicated in S4D) and re-induced with dox (200 ng/ml). Lysates were probed with antibodies specific for Rab25 or Tubulin. Black arrow indicates endogenous Rab25 in OVCAR-3 cells. F) Spinning disc confocal images of mNeonGreen-Rab25 DExCon cells (Rab25, green; Teton3G, blue), re-sorted as indicated in S4D. Scale bar=100 μm. G) Schematic illustration of the photoactivatable split PA-Tet-OFF and PA-Tet-ON constructs^14^ combined with DExCon (LUXon). H) Spinning disc confocal images (20x (left) or 63x (right) objectives) of mNeonGreen-Rab25 LUXon cells (Rab25 in green) expressing PA-Tet-OFF with mCherry-NLS reporter (red) 18h after being illuminated by blue light (10 hours) with or without dox treatment. Orthogonal views of top (1) or two bottom cells (2) are also shown with kymograph corresponding to white dashed line (1s interval, total 1 min). Scale bar: 10 μm. See movie S5-S6. I) Schematic of 3D cell-zone exclusion invasion assay. 1-4 indicate zones with different light illumination intensities across same well and these zones are also indicated in J): mNeonGreen-Rab25 LUXon cells (Rab25 in blue) expressing PA-Tet-OFF with mCherry-NLS as nuclear reporter (red) migrated for 24 h into cell-free collagen matrix labelled by FN-647 (green) while being illuminated by blue light of varying intensity. Dashed line indicates scratch and white arrows indicate the most invasive cells. Scale bar=100μm. See movie S7. K) Spinning disc confocal images of spheroids formed by mNeonGreen-Rab25 LUXon (PA-Tet-OFF2) cells illuminated with blue light for different times across 2 days as indicated (total 4h vs 21 hours). Cells invading collagen matrix supplemented with FN-647 (magenta) were labelled with Hoechst 3342 (blue) for 1 h prior imaging. Merge of all three channels, white rectangles for the mNeonGreen (Rab25, green) channel only, or zoom of the yellow rectangle is shown. Scale bar=100μm. L) Schematic of spheroid invasion assay “on chip” and illumination protocol used in M). mNeonGreen-Rab25 LUXon (PA-Tet-OFF2) cells (scale bar=1mm) together with N) quantification (n = 20-32 from 3 independent experiments, oneway ANOVA Tukey post hoc test). All Schematic illustrations were created with BioRender.com.

We next explored tunability of Rab25 expression. Exposing cells to different dox concentrations appeared to have a dose dependent effect when analysed by western blotting (S4C), however fluorescence microscopy revealed that the difference was primarily due to the number of responding cells (3D). Nevertheless, we noticed a positive correlation between expression of Tet-On 3G transactivator, reported as mTagBFP fluorescence, and mNeonGreen-Rab25 (S4D; R = 0.43, p = 0.0003). Taking advantage of this, we sorted for mNeonGreen-Rab25 expression and analysed Rab25 and mTagBFP levels. Dox-induced Rab25 expression levels in ‘low’ sorted cells were comparable to endogenous Rab25 expression levels in OVCAR-3 ovarian cancer cell lines over 48hours (3E) and Rab25 expression levels correlated with Tet-On 3G levels as expected (3F). The ability to re-induce previously sorted expression levels was preserved over several months under standard cell culture conditions, and reached maximal fluorescence after 72 hours (3E). These results indicate that the DExCon approach was able to re-activate endogenous silenced Rab25 on demand and levels could be tuned to match endogenous, physiologically relevant expression levels.

Whilst DExCon was able to temporally control Rab25 expression, this method lacks spatial control. We therefore explored the newly developed photoactivatable (PA)-Tet-OFF/ON system^14^, which has not previously been applied at an endogenous locus, to establish a light inducible ‘LUXon’ variant of DExCon. This system integrates the cryptochrome 2 (Cry2) cryptochrome-interacting basic helix-loop-helix 1 (CIB1) light-inducible binding switch with Tet-binding domains of Tet repressor (TetR) (residues 1–206) as the split DNA-binding domain, and the transcription activation domain of p65 (p65 AD) (3G). We tested PA-Tet-OFF or PA-Tet-ON and versions with improved blue light sensitivity (ON2; OFF2), stably delivered by lentivirus. Cells stable expressing PA-Tet-ON2, PA-Tet-OFF1 or 2 were then edited by mNeonGreen DExCon knock in to the Rab25 locus, illuminated by blue light for 24 hours and sorted, all showing the expected Rab25 endosomal localization (3H; S4E-G). Cyclical exposure to light/dark led to repeatable resilencing of Rab25, and the level of re-activation could be modulated by timing of blue light which is one of the major advantages of a lightinducible gene expression system (S4E-G). As these systems responded to the weak blue light exposure used (~6.8 W/m^2^), we found the additional layer of dox control to prevent leakiness of the system very practical. Dox addition with blue light was necessary to reactivate Rab25 in PA-Tet-ON2 system (S4E), while dox addition to PA-Tet-OFF1/2 completely blocked expression (S4F-G). Interestingly, in addition to the expected Rab25 localization, we observed some Rab25 positive vesicles in close proximity of the nucleus. These vesicles were significantly less mobile compared to others suggesting that the nucleus could under some circumstances sterically block movement of vesicles (3H; movie S5-6).

To demonstrate local induction of gene expression from the endogenous locus, we utilized a modified 3D cell-zone exclusion assay^23^. In this assay, a monolayer of the cells grown on a thick layer of collagen supplemented with non-labelled fibronectin is wounded and then overlaid with another layer of collagen supplemented with far-red labelled fibronectin (3I). The advantage of this approach is that system can be partly illuminated and Rab25 expressing cells can be monitored migrating towards the far-red fibronectin labelled cell-free area. This approach allowed us to demonstrate that Rab25 expression can be induced locally in 3D matrices and suggested a link between Rab25 re-expression and increased invasiveness (3J; movie S7). To quantify this, we next illuminated cell spheroids overlaid by 3D matrix, modulating the exposure to blue light (3K-N). Longer exposure to blue light led to brighter mNeonGreen signal in both PA-Tet-OFF/ON systems with expected Rab25 localization in polarized/elongated cells interacting with matrix fibrils (3K; S4H). When all cells were preilluminated 2 days before invasion to induce Rab25 expression, we saw a significant dose dependent effect of Rab25 induction via light exposure for over 6 hours on invasion that was not observed in the presence of dox in the TET-OFF system (3L-N). Adjusting the illumination protocol (S4I), activating Rab25 expression with constant blue light for 24hrs before spheroid formation/invasion, led to a light-dose dependent effect where the most invasive cells were those illuminated the longest approach (S4J-K). This demonstrates that tight control of gene expression levels and their impact on complex cell behaviour can be easily achieved/studied by the LUXon system.

### Triple knock-in determines Rab11 family colocalization

After generating mNeonGreen-Rab11a/ mCherry-Rab11b double knock in cells, we challenged ourselves to generate triple knock in A2780 cells (Rab11a/Rab11b/Rab25; 4A). To do so, mNeonGreen-Rab11a / mCherry-Rab11b double knock in cells were further genetically manipulated to introduce mTagBFP2 as part of the DExCon module controlling Rab25 expression. Dox treatment successfully specifically re-activated Rab25, visible as mTagBFP2, on the background of double knock in mNeonGreen-Rab11a/mCherry-Rab11b cells (S5A, B). Interestingly, activation of Rab25 expression increased the proportion of Rab11a and Rab11b localization at ERC spots, suggesting that Rab11a/b/25 networks are interconnected and can be modulated by Rab25 expression (S5C). Because mTagBFP2-Rab25 DExCon’s fluorescence intensity was less intense than that of mNeonGreen-Rab25 DExCon and Rab11a/b knock ins, triple knock in cells were re-sorted for bright mTagBFP2 signal and dox treatment kept over 94 hours prior fluorescence microscopy highlighting the tunability of expression with DExCon (S5A, B).

**Figure 4.**
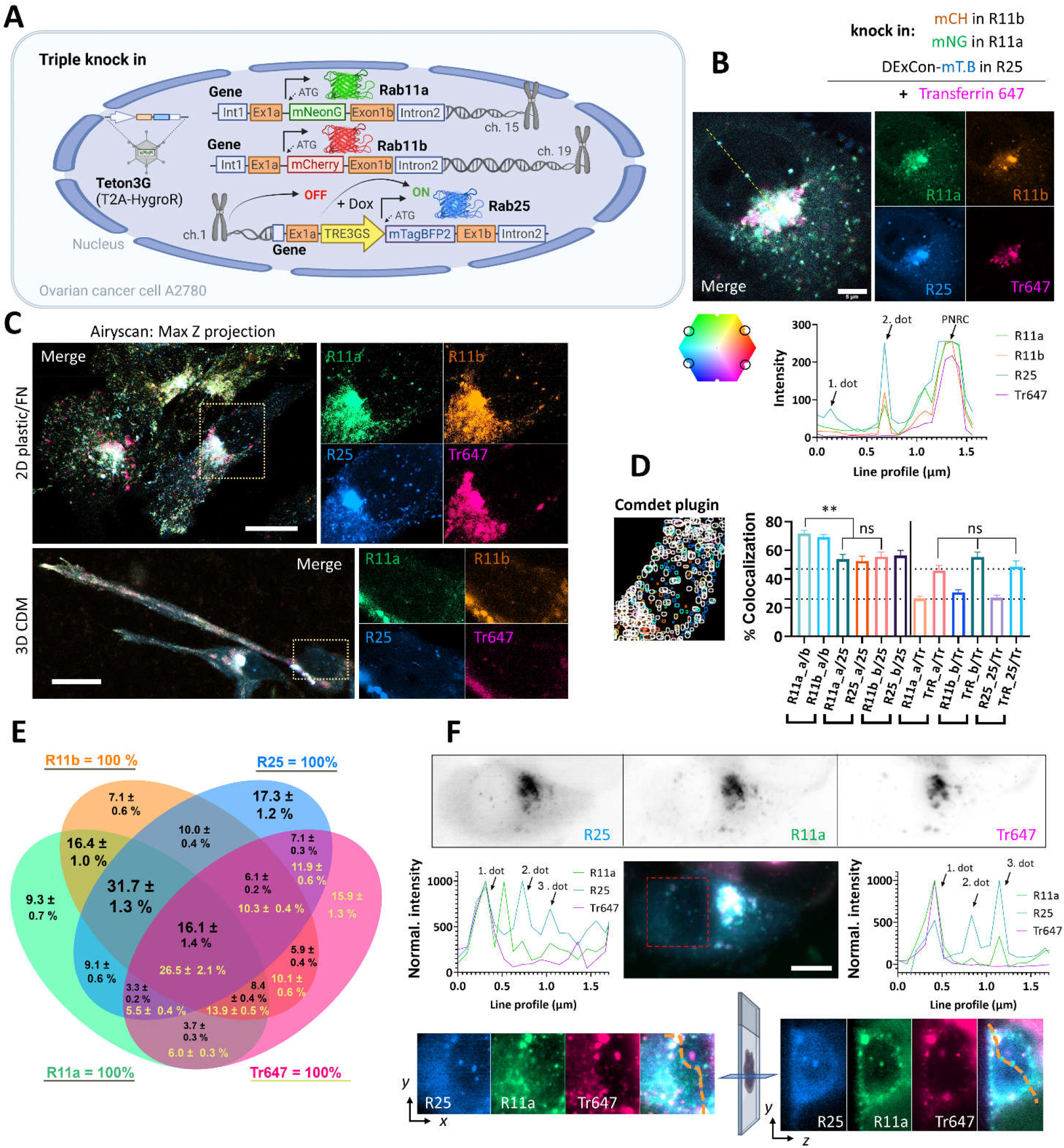
Simultaneous visualisation of Rab11 family members. A) Schematic of triple knock in A2780 cells (mNeonGreen-Rab11a; mCherry-Rab11b; DExCon mTagBFP2-Rab25); created with BioRender.com. B-D) mNeonGreen-Rab11a/mCherry-Rab11b knock-in A2708 cells were further modified with a DExCon-mTag-BFP2 module at the Rab25 locus (Tet-On transactivator introduced by lentivirus with hygromycin selection). Airyscan confocal fluorescence images of triple knock-in cells treated by dox (>94 h) trafficking Alexa-647 labelled Transferrin (Tr647; 15-60 minutes). Colours represent Rab11s as indicated and line profile corresponds to yellow dashed line. B) Scale bar=5μm. C) Maximum intensity Z-projections: top (2D, FN-coated), bottom (3D-CDM). Scale bar=20μm; see also movie S9-10. D-E) dox induced (>94h) triple knock-in A2780 cells recycling Tr647 were imaged and mNeonGreen/ mCherry/mTagBFP2/Alexa-647 positive vesicles tracked using Comdet plugin. D) Bars represent the percentage of colocalizing vesicles (100 cells, 46 000 vesicles (Rab11s); 23 000 vesicles (Transferrin)) and the contribution of individual channels (25- Rab25; Tr- Transferrin; a- Rab11a; b- Rab11b). One way ANOVA Tukey post hoc test used for statistical analysis. E) Venn diagram with the percentage of colocalization for every channel as 100% total; black numbers for Rab11s; yellow for Transferrin. F) Lattice light-sheet imaging of dox induced (>94 h) triple knock in A2780 cells recycling Tr647 are shown in grey as individual channels (one focal plane) or as Maximum Intensity Z-projection for merged channels; scale bar=10μm. Line profiles (normalized to maximal intensity, background subtracted) correspond to orange dashed lines, which connects the same vesicles (1. and 3. dot) in the 3D cell volume. See also S5E-F and movie S11-12.

Rab11 family members have been reported to colocalize with transferrin receptor and control its recycling^24,25^. To compare the colocalization of Rab11 family members and their potential contribution to the transferrin receptor recycling, we allowed cells to internalize Alexa-647 labelled transferrin (TFN) and analysed colocalization using AiryScan confocal and lattice light sheet microscopy (4B-F; S5B-E; movie S8-12). Whilst there was clear colocalization between Rab11a/b/25 and Tr647, some vesicles were highly enriched for Rab25 compared to Rab11a/b, but lacked TFN. Rab25 enriched vesicles negative for TFN were found in close proximity to the nucleus in approximately 90% of triple knock in cells (4B), both in 2D and 3D environments (4C; movie S9-12). To quantify the colocalization of vesicles we tracked in total around 46 000 vesicles per Rab11 family member. This analysis revealed that there is no significant difference in TFN colocalization with Rab11 family members, but Rab11a colocalizes with Rab11b to a significantly higher extent than with Rab25, or Rab11b with Rab25, (4D). Rab11a/b/25 largely colocalize with TFN, one another or all at the same time (16.1 % of all Rab11 vesicles/26.5 % of TFN vesicles; 4E) and that differences between Rab11 family members are mainly due to the unique Rab25 recruitment to a subset of vesicles (4F; S5D-E).

These results demonstrate that DExCon is a tool well suited for microscopy that allows multiple gene modifications simultaneously, including reactivation of previously silenced genes. Furthermore, complete integration can be verified using fluorescence-based sorting in polyclonal populations, and the approach provides a relatively simple platform for conditional rescues/knockouts that are, in contrast to FLP/Cre-driven recombination, reversible.

### Protein expression kinetics, stability and dynamics revealed using DExCon

Intrigued by slow expression kinetics of reactivated mNeonGreen-Rab25 DExCon in A2780 cells, we generated Rab11a and Rab11b DExCon cells to compare kinetics. Western blot suggested that we can detect mNeonGreen-Rab11a from 4h and mCherry-Rab11b from 6h after dox induction, while Rab25 DExCon was not detected before 8h (5A). The rate of dox induced expression was then quantified on individual DExCon Rab11 family gene modifications. mNeonGreen-Rab11a DExCon reached ~75 % of its maximal expression peak at 24h, significantly faster than other family members: at 24h mCherry-Rab11b DExCon reached at ~50 % peak expression, whereas mNeonGreen-Rab25 DExCon only reached ~40 % (5B). Similar results were also obtained from nanobody-mCherry-DExCon modifications (S6A-B) suggesting that observed differences can be explained by differences in the regulation of Rab11 gene expression and not by different FPs.

**Figure 5.**
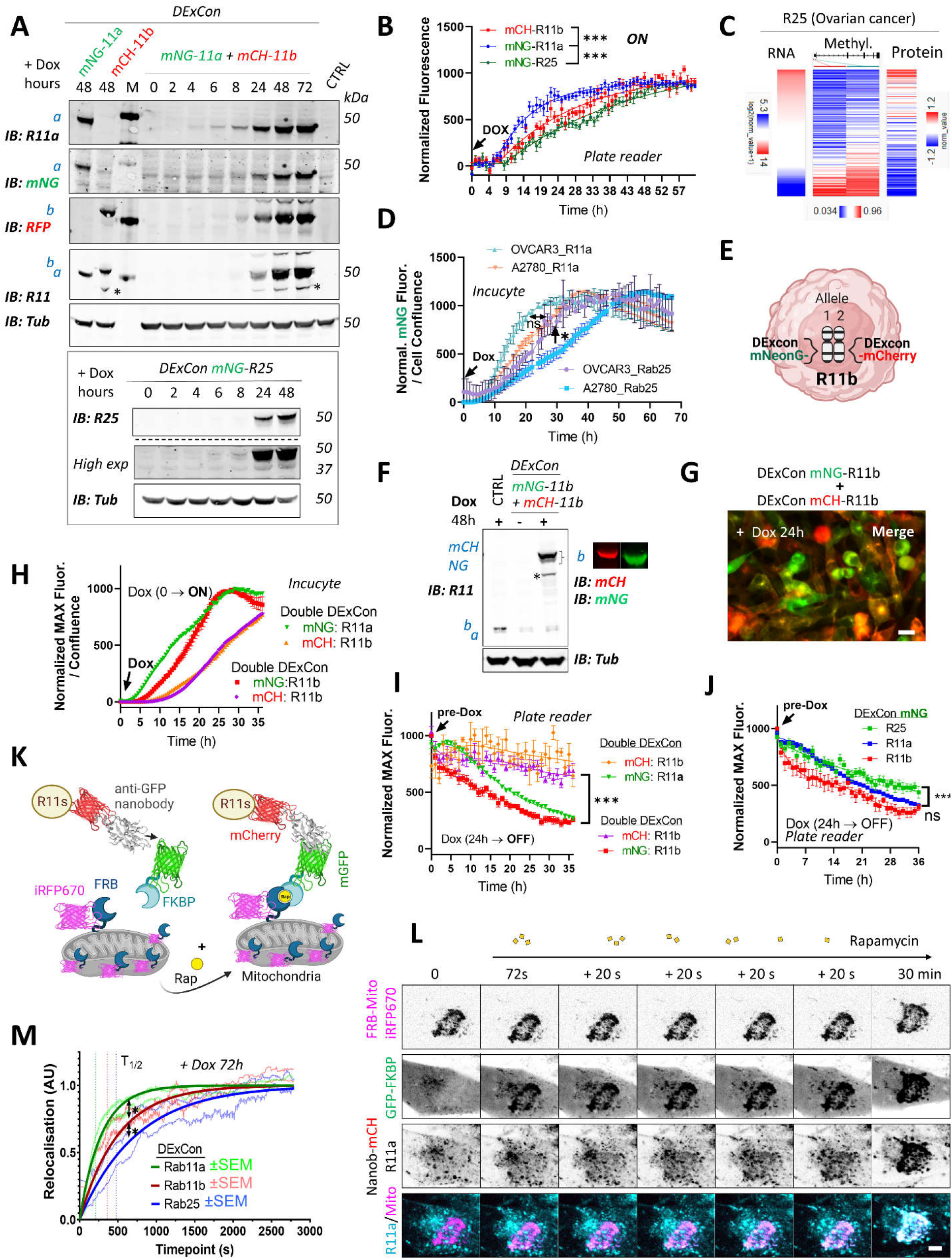
DExCon reveals protein expression kinetics and dynamics of relocalisation. A) Immunoblots of <u>mN</u>eon<u>G</u>reen-Rab11a and <u>mCh</u>erry-Rab11b DExCon or double DExCon A2780 cells (TOP) and mNeonGreen-Rab25 DExCon cells (Bottom) treated or not treated with dox for indicated amount of time (hours). Anti-mNeonGreen (mNG), anti-mCherry (RFP; mCH), anti-Rab25, anti-Rab11a or anti-Rab11 targeting both Rab11a/b (Rab11) were used to probe expression levels. Tubulin (Tub), loading control. M= marker; CTRL = un-modified wt A2780; a = Rab11a, b = Rab11b, c = Rab25 ± fluorophore knock in. Stars indicate mCherry/iRFP670 lower molecular weight band caused by hydrolysis during sample preparation^22^. B) Comparison of expression kinetics of DExCon-modified mNeonGreen-Rab11a/mCherry-Rab11b/mNeonGreen-Rab25 (dox added at time 0; arrow) using a microplate reader BioTek Synergy H1. Cells were seeded confluent and fluorescence normalized to the maximal intensity. One way ANOVA and Holm-Sidak post hoc tests were used for statistical analysis: n=6-9 from 3 independent experiments comparing normalised fluorescence intensity at 24h. C) Bioinformatic analysis of DNA promoter methylation (Methylation27K) of Rab25 and its expression (RNAseq; RPPA) in ovarian cancer (OV) using UCSC Xena (TCGA Pan-Cancer study). D) Comparison of expression kinetics of DExCon modified mNeonGreen-Rab11a or -Rab25, generated in A2780 or OVCAR-3 cells, induced by dox at time 0 (left arrow) and analysed using Incucyte® S3 imaging system. Readings were normalized to maximal fluorescence intensity and cell confluence (representative graph is shown with mean ± SD; one way ANOVA and Tukey post hoc test used for statistical analysis). E) Schematic of double DExCon modification, targeting both Rab11b alleles/loci with different fluorophores within the same cell. F) Immunoblots or G) fluorescence images (scale bar=100μm) of mNeonGreen/mCherry-Rab11b double DExCon cells A2780 treated ± dox. Blots were probed with anti-mNeonGreen (mNG), anti-mCherry (RFP), anti-Rab11 targeting both Rab11a/b (Rab11) specific antibodies. Tubulin (Tub), loading control. CTRL = un-modified wt A2780; a = Rab11a; b = Rab11b ± knock-in fluorophore. H-I) Comparison of mNeonGreen-Rab11a/mCherry-Rab11b or mNeonGreen-Rab11b/mCherry-Rab11b double DExCon expression kinetics H) triggered by dox (arrow; normalized and shown as in D) or their protein stability I). J) Comparison of DExCon induced mNeonGreen-Rab11a/Rab11b/Rab25 protein stability. I-J) Cells pre-induced with dox for 24h followed by dox removal (see arrow; microplate reader BioTek Synergy H1). Cells were seeded confluent and fluorescence normalized to the maximal intensity. Mean±SEM (9-18 repeats gathered from 3-6 independent experiments). Curves were fitted by fifth order polynomial function; one way ANOVA with Tukey post hoc test used for statistical analysis. K) Schematic of knocksideways combined with DExCon antiGFPnanobody-mCherry-Rab11 family modification. L) Representative confocal fluorescence timelapse images of antiGFPnanobody-mCherry-R11a DExCon modified cells induced with dox for 72 h (scale bar=5μm; see S7A-B for Rab11b and Rab25). Cells were co-magnetofected with FRB-Mito-iRFP670 and GFP-FKBP on 96 well ibidi imaging plate 24 hours before imaging and their heterodimerization induced by 200 nM Rapamycin as indicated. Halftime of Rab11a/b/25 relocalisation fitted and quantified M); for statistics see S7C, n = 10-18 from 3-4 independent experiments.

Since Rab25 is silenced in A2780, we performed bioinformatic analysis of its methylation status. We found that there is an inverse relationship between methylation of Rab25 promoter and its expression, where decreased methylation is correlated with increased gene expression, at both mRNA and protein levels across all cancer types (5C; S6C). ATAC-seq dataset analysis further suggested that the chromatin structure of the silenced Rab25 locus, both in promoter and enhancer sequences, is less accessible for transcriptional machinery to drive transcription (S6C). To prove that slower Rab25 DExCon expression kinetics in A2780 cells can be simply explained by the decrease of DNA accessibility for transcription, we knocked in TRE3GS-mNeonGreen to the Rab11a or Rab25 loci of OVCAR-3 cells, an ovarian cancer line where both Rab11a and Rab25 are endogenously expressed. Dox induction again led to the correct localization of Rab11a or Rab25, as expected (S6D), but this time there was no delay in the normalized expression kinetics of Rab25, as was observed for A2780 (5D). Interestingly, in both A2780 and OVCAR-3 cells Rab11a DExCon always led to greater fluorescence intensity compared to Rab25 DExCon, suggesting additional mechanisms of protein level control (S6E).

We further explored the possibility of simultaneous visualization of different alleles of the same gene (5E). We were able to successfully modify the second allele of Rab11b by knocking in TRE3GS-mNeonGreen sequentially into mCherry-Rab11b DExCon cells. Cells were sorted for highly homogenous expression levels of mNeonGreen/mCherry upon 48 h of dox induction (S6F) and dox-mediated re-induction (24-48h) led to activation of both alleles (5F) with consistent localization (5G). Unexpectedly, expression kinetics were noticeably different between mCherry and mNeonGreen modified Rab11b (5H; movie S13). While the mNeonGreen kinetics of Rab11a was still slightly faster than mNeonGreen of Rab11b, the mCherry Rab11b fluorescence displayed significant delay similar in both Rab11a/b and Rab11b double DExCons cells. When DExCon cells were pre-induced with dox for 24 hours, followed by dox removal, a decrease in fluorescence was observed over time (5I). mCherry-Rab11b showed remarkably stable fluorescence, whereas mNeonGreen-Rab11b was lost more rapidly. Stable fluorescence of mCherry was also observed for nanobody-mCherry DExCon Rab11s (S6G). By contrast, mNeonGreen fluorescence continuously declined overtime, where Rab11a and Rab11b decreased in intensity more rapidly than Rab25 following dox removal (5J). This suggests that the FP is an important consideration when using FP-knock-in to understand protein dynamics, but opens the possibility of modifying individual (e.g. mutant versus non-mutant) alleles within cells to study their stability and function.

The localisation of Rab GTPases is fundamental to their function, but the relative stability of Rab11 family members at their endogenous location is not known. We therefore combined DExCon with knock-sideways, a method of protein relocalisation within live cells^26^, to give insight into the dynamics of Rab11a, Rab11b and Rab25 localisation. Relocalisation to mitochondria was achieved by rapamycin (rap) induced dimerization of FKBP-mGFP with mitochondrially targeted mito-FRB-iRFP670 in GFPnanobody-mCherry DExCon modified A2780 cells (5K) and the dynamics of Rab11 family members analysed by time-lapse imaging (5L; S7A-B). Direct comparison of Rab11a/b/25 dynamics revealed an ultra-fast relocalisation of Rab11a (t_1/2_=209s), significantly slower Rab11b relocalisation (t_1/2_=362s) and slower still dynamics of Rab25 (t_1/2_=476s) (5M; S7C).

Taken together these results show our DExCon approach can be used to reveal subtle but significant differences in protein expression kinetics and protein dynamics, and that despite their similarity at the protein level, such differences are observed within the Rab11 family.

### DExogron: a tunable approach to modify protein levels within cells

Because removal of dox did not rapidly decrease DExCon-driven protein levels, we added a second layer of control utilizing the improved Auxin-inducible degron (AID) system reported with low basal degradation^27^. We combined our DExCon module with an auxin-inducible destabilizing domain (short degron miniIAA7) that allows proteasomal degradation in an auxindependent manner to generate DExogron gene modifications (6A).

**Figure 6.**
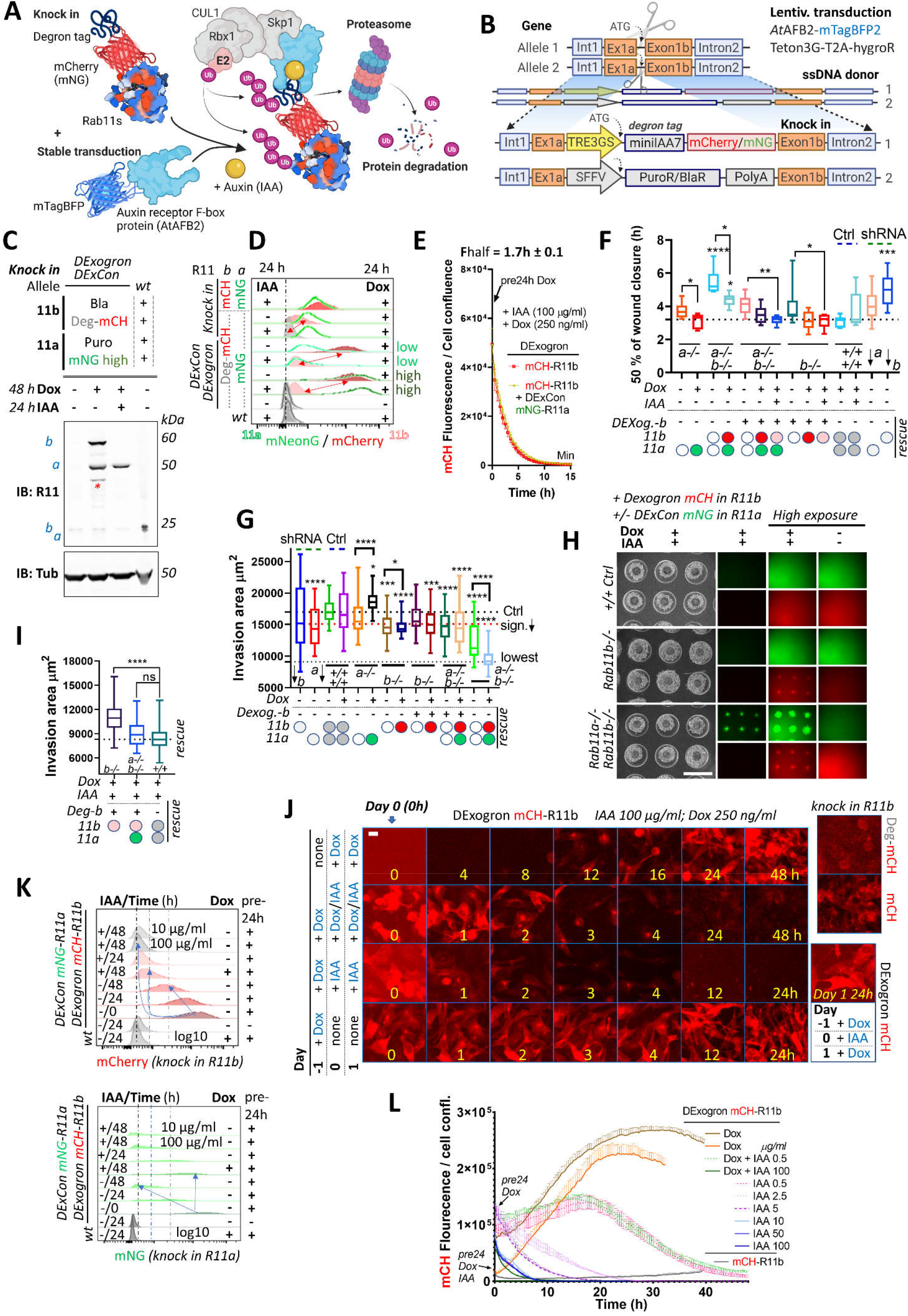
DExogron: a tunable approach to modify protein levels within cells. A) Schematic illustration of an improved AID system with low basal degradation^27^ used in this study. Degron tag=miniIAA7 B) Schematic of components of knock in cassette used for DExogron, together with second donor cassette (providing antibiotic resistance) independently targeting another allele. A2780 stably expressing AtAFBP2 and Teto3G-T2A-HygroR were used for knock-in; mCH = mCherry as red; mNG = mNeonGreen as green. All Schematic illustrations were created with BioRender.com C) Immunoblots of <u>mN</u>eon<u>G</u>reen-Rab11a DExCon (sorted for high; the full blot with also cells sorted for low in S8F) / miniIAA7(=Deg)-<u>mC</u>herry-Rab11b DExogron cells treated or not treated with dox (250 ng/ml) ± IAA (Indole-3-acetic acid as a source of Auxin; 100 μg/ml). Bla/Puro represents <u>Puro</u>mycin or <u>Bla</u>sticidine resistance given by additional knock in outcomes. Blots were probed with anti-Rab11 antibody targeting both Rab11a/b (Rab11). Tubulin (Tub), loading control. wt= un-modified A2780; a = Rab11a; b = Rab11b ± fluorophore knock in. Stars indicate mCherry/iRFP670 lower molecular weight band caused by hydrolysis during sample preparation^22^. D) FACS analysis of DExogron-mCherry-Rab11b/DExCon-mNeonGreen-Rab11a (sorted for high or low) A2780 cells or classical R11a/Ra11b double knock ins cells ± miniIAA7 (=Deg) without the TRE3GS promoter are shown. Cells were treated ± dox (250 ng/ml) ± IAA (100 μg/ml) for 24h as indicated. Red arrows indicate the dynamic range of mCherry intensity change. wt= un-modified A2780. Dashed line indicates negative Ctrl. Log10 scale shown. E) Degradation kinetics of DExogron -mCherry-Rab11b (± DExCon-mNeonGreen-Rab11a; See also S8G) analysed by Incucyte® imaging. Cells were pre-treated by dox for 24h followed by dox/IAA co-treatment as indicated and half-time of mCherry fluorescence decrease calculated. The mean of 6 replicates from 3 independent experiments is shown±SEM. F) Confluent cells were pre-treated for 24 ± dox (250 ng/ml) ± IAA (100 μg/ml), scratch wounds introduced and imaged in phenol free RPMI±dox/IAA as indicated. a-/-:DExCon-mNeonGreen-Rab11a (rescue shown as green oval), b-/-: DExCon-mCherry-Rab11b (rescue shown as red oval) or Deg-b:DExogron-mCherry-Rab11b (pink oval indicates low mCherry expression). Box and whisker plot of 2D scratch wound migration (n = 3-9 repeats from 3 independent experiments), shown as time required to close 50% of the normalized wound area. One-way ANOVA analysis Tukey post hoc test used for statistical analysis. G-I) On-chip spheroid invasion assay of cells described in F), migrating (72 h) in collagen matrix supplemented with FN and treated ±dox (250 ng/ml) ±IAA (100 μg/ml) as indicated (pre-induced for 24h). G) Quantification of spheroid invasion ±dox (n = 48-76 from 3 independent experiments). Black dotted line indicates Ctrl, red line is the threshold of statistical significance (see right arrow; statistics as described in F). See also S9D-E. H) Brightfield and fluorescence images of spheroids with lower or higher exposures (scale bar=1mm) co-treated by ±dox/IAA and quantification of their invasion capacity I). J) Timelapse images of DExogron-mCherry-Rab11b A2780 cells (Incucyte, scale bar=20μm). Cells were treated with dox/IAA as indicated. mCherry fluorescence intensity compared with classical endogenous (Deg)-mCherry-Rab11b tagging (on the right). See movie S14. K) FACS of DExCon/DExogron-R11a/R11b cells treated as indicated (dox 250 ng/ml; IAA 100 μg/ml if not stated otherwise). Pre-24 h = dox pre-treatment for 24 h prior analysis; wt= A2780 without CRISPR knock in. Blue arrows indicate the rate at which signal intensity decreases. Log10 scale is shown. L) The effect of different dox, dox/IAA or IAA levels (as indicated cells were dox or dox/IAA pre-treated for 24h) on fluorescence of DExogron-mCherry-Rab11b (conventional (deg)-mCherry-Rab11b knock-in for comparison without any treatment; top right) analysed by Incucyte® S3 system from images as shown in L). Representative curves of mCherry fluorescence normalized to cell confluence (3 replicates ± SEM) are shown.

Because we were able to independently target both alleles of Rab11a/b, we first combined our knock-in strategy with knock-in of a cassette composed of puromycin (PuroR) or blasticidine (BlaR) resistance markers and PolyA signal to block endogenous expression (6B; S8A). In A2780 cells, inclusion of an additional promoter sequence (SFFV) was required because endogenous promoter-driven expression was insufficient to yield antibiotic resistance, in contrast to HEK293T cells. Sequential targeting of the second Rab11a or Rab11b allele in DExCon cells using PuroR or BlaR, significantly improved knock in outcome both in DExogron, DExCon Rab11b or double DExCon Rab11a/b cells (S8B-D) and thus successfully generated complete knock in/knock out cells with only one allele of Ra11a or b under dox control (rescue). Notably, simultaneous targeting of two Rab11a alleles by DExCon and PuroR dramatically increased the proportion of DExCon knocked in cells surviving puromycin treatment, from 0.2 to 25 % (S8E). We successfully generated combinations of DExCon/DExogron-Rab11a/b, with additional PuroR/BlaR knock-ins (6C; S8F). Dox treatment led to rescue of protein expression as expected and simultaneous addition of Indole-3-acetic acid (IAA as a source of Auxin) efficiently depleted dox-induced expression only when the degron tag was present (DExogron minilAA7-mCherry-Rab11b) (6C; S8D, F; movie S14-15). Flow cytometry analysis confirmed that IAA/dox co-treatment (24-48 h) decreases DExogron mCherry-Rab11b fluorescence to the level of un-treated Rab11b miniIAA7-mCherry knock in cells with degradation half time kinetics of 1.7 ± 0.1 h (6D, E; S8G; movie S14).

Introducing a degron tag using one donor without additional DExCon modification led to a decrease in mCherry fluorescence (6D) and revealed two additional drawbacks. A significant proportion of cells did not respond to IAA likely due to the truncation of the minilAA7 sequence during HDR (S8H). This could be overcome by including negative sort after treatment by IAA in our protocol (see diagram S8I). However, treatment with IAA resulted in 90≥70% mCherry removal, which was close to the control level but not complete, limiting the effectiveness as complete removal tool (6D).

Rab11 family GTPases are implicated in cell migration, invasion and cancer progression ^10,15,28–30^. To test the functionality of DExogrons we performed a 2D scratch wound migration assay and found that Rab11a or Rab11b DExogron modified cells closes the wound significantly later than control wt cells in the absence of dox induced expression, and this phenotype is further amplified by Rab11a/b knockout combinations (6F; S9A; movie S16). Knockout phenotypes were reversed when Rab11a/b expression was reactivated with dox, however induction of degradation with IAA did not oppose this rescue, suggesting that IAA induced Rab11b depletion is incomplete in the presence of dox and is not sufficient to mimic the knockout phenotype. The importance of Rab11a or Rab11b for cell migration was further verified with A2780 stably expressing shRNA anti Rab11a or b (6F; S9A-B). The effect of proliferation on wound healing closure was negligible, as any change in Rab11a/b expression did not alter cell growth (S9C). Notably, we observed a toxic effect of IAA (100 μg/ml) from 10 h of exposure when A2780 cells seeded sparsely.

To investigate how manipulation of Rab11a/b expression influences cell migration/invasion into 3D collagen/FN matrix, we performed spheroid invasion assays. Reducing Rab11a or b expression using shRNA, DExCon, DExogron or their combination decreased the ability of cells/spheroids to invade FN-rich collagen and this was significantly rescued by dox treatment in Rab11a DExCon single knock in cells (6G; S9D-E). Dox-induced re-expression of Rab11b had a dose dependent dominant negative effect on invasion under all combinations, sufficiently blocking even Rab11a rescue. The negative effect of Rab11b rescue was reversed by simultaneous dox/IAA co-treatment and led to cell invasion comparable to wt control cells or even significantly higher (6H-I) despite the cytotoxic effect of IAA observed on individually migrating cells (S9D-E vs 6H). This suggests that cells are sensitive to Rab11b re-expression levels and that balanced dosage is needed for coordination of membrane trafficking important for 3D cell migration.

We next tested whether induction of degradation with IAA can improve the AID pitfall of incomplete Rab11b depletion by withdrawing dox following re-activation of Rab11b expression. Indeed, mCherry fluorescence intensity of DExogron Rab11b cells or DExogron-Rab11b/DExCon-Rab11a cells was slightly, but significantly, decreased below the fluorescence of dox/IAA co-treated cells as early as 1-4h after switching to dox-free media supplemented only with IAA (6J; S10A) and returned to nondetectable levels in 24-48h (6K). After 48h, expression of DExCon mNeonGreen-Rab11a was reduced, but still detectable (6K). These data show that the DExogron is an important modification that can lead to complete removal of protein following withdrawal of dox-induced gene expression.

Because IAA appeared to influence cell proliferation, we dropped from 100 μg/ml to 0.5 μg/ml and found that degradation was slowed, but still occurred to a similar extent. Dox/IAA co-treatment was sufficient to tune DExogron mCherry-Rab11b to endogenous levels (6K, L; S10B). DExogron-Rab11a / DExCon-Rab11b similarly resulted in tuned steady-state fluorescence intensity of mNeonGreen Rab11a with degradation half time kinetics around 2.5 ± 0.1 h (S11A-E; movie S17).

These data therefore indicate that DExogron can eliminate the shortcomings of the AID system, allowing fine-tuning of expression levels up to complete depletion following CRISPR based modification. Furthermore, our results suggest that although Rab11b and Rab11a may have a complementary function in migration and invasion of A2780 ovarian cancer cells, they differ substantially in their post-transcriptional regulation.

## Discussion

Here we describe DExCon, LUXon and DExogron tools to reversibly modulate gene expression from the endogenous locus. These systems allow effective knockout of gene expression, but allow subsequent re-activation of transcription (or activation of transcriptionally silent genes). LUXon permits the spatial and temporal control of gene expression, and DExogron adds a further level of control such that expression levels can be tuned appropriately, or complete degradation induced.

Significant effort has been made to achieve conditional control of gene expression at the endogenous locus in order interrogate gene function in a native biological and pathological context. Binary recombination approaches, such as the Cre/loxP system, allows inducible inactivation^2^, or activation through conditional excision of a stop element in the target gene^31^. However, the ability to control gene expression is lost following irreversible deletion, and this system does not allow evaluation of the phenotypic effects across a spectrum of different expression levels. Our DExCon/Dexogron /LUXon approaches are attractive alternatives as they offer, in addition to reversible inactivation of protein coding genes, conditional and (for LUXon) spatiotemporal expression control of reactivated genes which are visualized by an introduced FP.

Other approaches to modulate endogenous gene expression have used components of lac repression and tet activation systems introduced to the endogenous promoter sequence or downstream, including “Reversible Manipulation of Transcription at Endogenous loci (REMOTE)-control”^32^. Our approach differs because it bypasses the function of the endogenous promoter and does not require insertion of effector binding sites that can perturb target gene expression. Our one-CRISPR knock in based method is best suited for any genes whose protein-coding transcripts share the initial ATG codon and can be N-terminally tagged, or introduce the DExCon module upstream of alternatively edited genes. For genes where the fluorophore would have to be introduced internally or C-terminally, two CRISPR knockin step approaches are necessary (one for TRE3GS promoter, second for FP). Although our system does not silence transcription driven by an endogenous active promoter, it generates nonsense frameshifts in transcripts originating from the endogenous promoter and can activate silenced transcripts. This offers advantages over methods that alter promoter methylation (e.g. dCas9 fused to transcriptional activators or repressors^33^, which can create modifications over several tens of kilobases and impact off-target genes^34,35^) and overcomes the limit of classical knock-in where low expression confounds detection or protein products. Furthermore, in contrast to transient transfection of exogenous vectors, DExCon preserves splicing control and allows analysis of the full complement of physiologically localised isoforms within cells.

The AID system is an excellent method of protein down-regulation^7^, which has since been further developed^8,27,36^ allowing rapid degradation of target proteins. Using CRISPR we successfully applied the improved AID system to the endogenous locus of Rab11b^27^, however the incorporation of degron miniIAA7 tag in front of mCherry at the Rab11b locus resulted in a profound decrease in mCherry fluorescence without induction by IAA, and induction did not lead to complete protein removal (6D). Our combination of degron and DExCon (DExogron) however can generate robust expression and allows additional fine temporal control of DExCon system, providing a system that can reveal acute phenotypes that otherwise might be masked by compensation or a dominant terminal phenotype.

For broad application of strategies such as DExCon, feasibility and efficiency are critical. One of the main obstacles to gene editing via HDR is its low efficiency, which complicates targeting all alleles, generation and selection of homozygous knock-in modifications^4^. Our optimized magnetofection mediated delivery of high fidelity Cas9 as a ribonucleoprotein complex with single-stranded donor DNA ensured precise and on-target integrations to endogenous Rab11 family gene loci reflected by expected localization in cells (1C). Although truncated integration events are often reported^21^, functional flanking components (tet-responsive promoter and FP) allows selection of polyclonal populations with full integration of the ssDNA template (including for example the nanobody anti-GFP domain), negating the need for comprehensive and labour-intensive genotyping (2H-I). We demonstrated the robustness of our approach by generating single, double and triple knock in DExCon at the endogenous loci of Rab11 family genes in A2780 without the need for clonal selection. Our strategy further offers sufficient expression modulation from monoallelic modifications, therefore we could independently target alleles of the same gene using combinations of different donors (DExCon-mNeonGreen, DExCon-mCherry, antibiotic resistance (5F; S8B-F)). As proof of principle, we imaged protein products of both Rab11b alleles to demonstrate new possibilities, for example studying the dynamics of mutant and wildtype alleles of oncogenes and tumour suppressors. When simultaneously delivering DExCon together with antibiotic resistance to a second allele, we observed a dramatic increase in selection efficiency from 0.2 to 25% (FP-positive dox-responsive cells surviving antibiotic treatment) allowing for easy enrichment of homozygous edits (S8E). Sequential delivery of antibiotic resistance to DExCon-Rab11a and -Rab11b (S8B-F) further improved efficiency, effectively generating a KO of one allele to ensure that all protein generated by cells is tagged appropriately (mNeonGreen/ mCherry in this case), full knockout with dox-controlled expression. The conventional Tet-OFF/ON system is widely used in various research models, such as transgenic animals^37^, and can be easily combined with our DExCon/DExogron/LUXon strategy for a huge range of applications across the field of biology. Notably, DExogron harbours promising application also for essential genes with knockout lethal phenotypes. Here, the continual presence of dox or the transition from Tet-on to Tet-off system would be required, and IAA treatment could induce immediate knockout effect.

To demonstrate the potency of our strategies on closely related genes, we deployed them to study members of Rab11 family (Rab11a, Rab11b, and Rab25) in an ovarian cancer cell line that lacks endogenous Rab25 expression (3A). We demonstrated that spatiotemporal re-activation of silenced endogenous Rab25 expression can promote invasive migration (3J-O), and that Rab11b and Rab11a may have complementary functions in migration and invasion (6F-I). In addition, our tools allowed us to quantify surprising differences between Rab11 family members. We found that our system reflects the accessibility for transcriptional machinery to drive transcription at the endogenous locus, demonstrating that Rab11a has significantly faster expression kinetics and compared to Rab11b and reactivated Rab25 expression is slower still (5A-C). In cells with endogenous Rab25 expression, expression kinetics are demonstrable faster however (5D). We also demonstrated that the ability to relocalise differed significantly between Rab11 family members with Rab25 as the least dynamic (5KM; S7A-C). Furthermore, we observed a population of Rab25 specific vesicles in the nuclear periphery which seems to be less mobile (3I) and these vesicles were always negative for transferrin receptor (4B-F). Such vesicles with limited mobility were recently described to be located in nuclear envelope where they exhibit propensity to undergo fusion and were described as a novel endosomal route from the cell surface to the nucleoplasm that facilitates the accumulation of extracellular and cell surface proteins (e.g. EGFR) in the nucleus ^38,39^. Rab25-mediated EGFR recycling^29,40^ and elevated levels of nuclear EGFR correlate with poor prognosis in ovarian cancer^41^, but since Rab25 has not been connected with nuclear envelope-associated endosomes (NAE) our findings warrant further investigation.

Overall, the work presented here opens new avenues not only for the precise expression control of endogenous genes, but also for the manipulation and imaging of gene function. All the benefits of gene editing can be obtained in one CRISPR based knock in. DExCon, DExogron, LUXon are powerful tools that allow new insight into the function of individual genes and closely related gene families.

## Methods

Detailed list of antibodies, chemicals, sequences, plasmids and resources used are described in supplementary excel file S1 or accessible from https://doi.org/10.48420/14999550.

### Constructs

A range of plasmid DNA constructs were used and generated in this study including 27 donors used for long ssDNA preparation, and these are listed with details in excel file S1. Plasmid maps are provided here https://doi.org/10.48420/14999526 (donors) and https://doi.org/10.48420/16810525 (others).

To prepare plasmid donors for ssDNA synthesis, first cDNAs coding for mNeonGreen/ mCherry / iRFP670 /anti-GFPnanobody/minilAA7/TRE3GS/SFFV-T2A-PuroR-polyA (and so on) or gene fragments (Rab11a/b/25) designed with universal linkers (including SpeI/AgeI or MluI site; see plasmid maps and catalogue of donors in excel file S1) were commercially synthesised by GENEWIZ (FragmentGENE), ThermoFisher (geneArt Gene Synthesis) or IDT (gBlocks) depending on the synthesis limitations (GC-rich/repetitive sequences; see excel file S1). These DNA sequences were delivered in pUC57 or pMKRQ vectors or cloned into pJET1.2 vector (CloneJET PCR Cloning Kit. #K1231; Thermofisher) and later combined using SpeI/AgeI/XhoI sites or by Gibson assembly. Gibson assembly and SpeI/AgeI/MluI/EcoRI sites were also used to add, switch or remove additional sequences (e.g. mTagBFP2 from pLenti Lifeact-mTagBFP2_PuroR). Lentiviral pCDH-antiGFPnanobody-mCherry-Rab11A (overexpressing control) was cloned using pCDH-EF1-tagBFP-T2A-mycBirA*-Rab11a (previously prepared in our lab) via SpeI/SalI and XbaI/XhoI complementary cleavage sites. pMito-mCherry-FRB^42^, a gift from Stephen Royle (Addgene plasmid # 59352), was used as a template for generation of pMito-iRFP670-FRB (used with GFP-FKBP for knocksideways). pSH-EFIRES-P-AtAFB2 (Addgene plasmid #129715)^27^, pCDH-EF1-tagBFP-T2A-mycBirA*-Rab11a and mTagBFP2 were used to generate lentiviral pCDH-AtAFB2-mTagBFP2. TRE3GS was removed by Cla/EcorI cleavage from 3rd generation tet inducible vector pCDH-TRE3GS-EF1a-tagBFP-T2A-TetOn3G (gift from Andrew Gilmore lab) to generate DExCon modules, and the plasmid modified by Gibson assembly to generate pCDH-EF1a-HygroR-T2A-TetOn3G with Hygromycin resistance (for DExogron). Other plasmids used (for details see excel file S1): pLenti Lifeact-iRFP670 BlastR^43^ (Addgene Plasmid #84385); pGIZ-PuroR-IRES-GFP_shRNA anti Rab11a/b (Dharmacon); GFP/mCherry-R11a/b (created by Davidson or Caswell lab); lentiviral plasmids coding two versions of PA-Tet-OFF/ON^14^. All constructs were verified first by restriction analysis and then by sequencing. Further details on the cloning strategy, and plasmids used in this study are available upon reasonable request or will be available at Addgene.

### Cell culture

A2780-DNA3^29^ and OVCAR-3 (ATCC) ovarian cancer cell lines were cultured in RPMI-1640 medium (R8758, Sigma). Ovarian cancer COV362 (ECACC), TIFs (Telomerase immortalised fibroblasts^29^) and human embryonic kidney (HEK)293Ts cells were cultured in Dulbecco’s Modified Eagles Medium (DMEM) (D5796, Sigma). All cell culture medium was supplemented with 10% v/v foetal bovine serum (FBS) and ciprofloxacin (0.01 mg/ml; Sigma) and cells were maintained at 37°C in a humidified atmosphere with 5% (v/v) CO_2_. Cells were confirmed as being negative for mycoplasma contamination by PCR. A2780 were selected for antibiotic resistance as follows: hygromycin 200 μg/ml; blasticidin 5 μg/ml; puromycin 0.5-1 μg/ml.

### Lentivirus packaging

Lentiviral particles were produced in HEK293T cells, via a polyethylenimine (PEI)-mediated transfection with packaging plasmids psPAX2 and pM2G with lentiviral vector pCDH or pLenti in the ratio 1.5:1:2. Similarly, retroviral pRetroQ was delivered with pM2G and specific gag-pol **plasmid**. Supernatants were collected at 94 h after transfection, filtered through a 0.45 μm filter and added to target cells. Transduced cells were selected by appropriate antibiotics or grown up and sorted by FACS.

### Flow cytometry and analysis

Cells (pre-induced ± dox ± IAA as needed) were gated based on mNeonGreen/ mCherry/ mtagBFP expression as desired and bulk sorted into 15 ml centrifuge tubes by FACS Aria II (BD). In few cases, cells were cloned by serial dilution. Sorted DExCon/DExogron cells pre-induced by dox were again screened for fluorescence activation after two weeks of growing in the asence of dox. Flow cytometry was performed using a LSR Fortessa (BD) with a five-laser system (355, 405, 488, 561, 640 nm) and fluorescence levels recorded with for 10,000 events with configuration as follows: mNeonGreen (488 530/30), mCherry (561 610/20), mtagBFP (405 450/50), and iRFP670 (640 670/14). For all experiments, cells were pre-treated ± dox ± IAA (24-72h) prior analysis and gated based on the empty A2780 (negative control). If analysis could not be done same day as cell preparation, cells were detached using PBS with 5 mM EDTA, fixed with 2% PFA on ice (10 min), and exchanged to 0.5 % BSA in PBS supplemented with 0.1% sodium azide. Data were processed using the FlowJo software (FlowJo LLC).

### Antibodies and reagents

The following antibodies and reagents were used (details in excel file S1): Rab11 (rabbit pAb, # PA5-31348), α-tubulin (mouse mAb DM1A, #ab7291), Rab11a (rabbit mAb D4P6P, #2413S), Rab25 (rabbit pAb, #13048S), RFP (rat mAb 5F8), mNeonGreen (mouse mAb 32F6-100). Fluorescent secondary antibodies (Li-cor or Invitrogen) were used as recommended by the manufacturer. The reagents used were blasticidin S HCl (Gibco) and puromycin dihydrochloride (Thermo), hygromycin B (Roche), fibronectin (sigma), high density type I collagen (Corning), SiR-Tubulin (Cytoskeleton), Dynabeads™ MyOne™ Streptavidin C1 (Thermo), SuperScript™ IV Reverse Transcriptase (ThermoFisher), and Agencourt AMPure XP (Beckman Coulter). Doxycycline hydrochloride (Sigma) was dissolved as 250 μg/ml in water (stored −20 °C in aliquiots) and used within 14 days if kept in 4 °C. Similarly, Indole-3-acetic acid sodium salt (source of Auxin; Cambridge Bioscience Limited, #16954-1g-CAY) aliquiots (10 mg/ml in water) were in stored −20 °C, but used within 2 days once melted.

#### SDS-PAGE and Quantitative Western Blotting

Cells were lysed in denaturing lysis buffer (2% SDS, 20% glycerol, 120mM tris pH 6.8, 0.1% bromophenol blue) and heated 10 min 98 °C. Cell lysates were resolved under denaturing conditions by SDS-PAGE (4–12% Bis-Tris gels; Invitrogen) and transferred to nitrocellulose membrane using Trans-Blot Turbo Transfer System (Bio-rad). Membranes were blocked with 4 % BSA-TBS (Tris-buffered saline) for 1h followed by incubation overnight at 4°C with the appropriate primary antibody in 2-3% BSA TBST (TBS with 0.05% Tween 20) and then incubated for 1 h with the appropriate fluorophore-conjugated secondary antibody in 2.5% milk-TBST. Membranes were scanned using an infrared imaging system (Odyssey; LI-COR Biosciences).

### Bioinformatic analysis

Rab11 expression levels (RNAseq, free of batch-effects) across different tissues and their relationship to cancer, were analysed using UCSC Xena **https://xenabrowser.net/** ^17^ from the TCGA TARGET GTEx study and graphs generated in Prism software or visualized using Qlucore Omics Explorer. UCSC Xena tool was used to compare the methylation pattern of Rab25 and its expression (TCGA Pan-Cancer or GDC Pan-Cancer study). Computed quantitative trait loci, eQTLs, of Rab11s isoforms across multiple healthy tissues (https://doi.org/10.48420/16988617) were visualized using The Genotype-Tissue Expression (GTEx) browser: **https://gtexportal.org/** (dbGaP accession number phs000424.vN.pN on 05/06/2021).

### Preparation of long ssDNA template

Single stranded DNA was prepared through optimized reverse-transcription of an RNA intermediate (IVT) for sequences up to 2 kb using high-fidelity Reverse Transcriptase, similarly to that described in detail in ^3^. Briefly, all donor plasmids (details in excel file S1) with a T7 promoter site followed by sequences used for knock in (fluorophore sequence flanked by 150-300bp homologous arms) were first linearized by the appropriate restriction enzyme (cutting just outside the homologous arm opposite to the T7 promote), followed by in vitro transcription (HiScribe T7 polymerase, NEB #E2040S), DNA degradation (TurboDNAse, Thermo #AM2238), reverse transcription (SuperScript™ IV, Thermo # 18090050) with RNAse Inhibitor (SUPERase, AM2696 Thermo) and RNA hydrolysis (95 °C 10 min in pH>10 with EDTA). Final ssDNA product and all intermediate steps were purified using SPRI beads (AMPure XP, Agencourt) according to manufacturer protocol. Longer ssDNA sequences (>2 kb) were then prepared using PCR with biotinylated forward primers^44^ (see diagram in excel file S1), final ssDNA purified using SPRI beads (AMPure XP, Agencourt) and anti-sense ssDNA strand used for knock in. Where donor plasmid was used as a DNA template, biotinylated PCR-product was purified using Dynabeads™ MyOne™ Streptavidin C1 (Cat# 65001, Thermo) and anti-sense ssDNA strand eluted by 20mM NaOH (later neutralized by HCl). All ssDNA were denaturated by heating (70 °C, 10 min) in RNA Loading Dye containing formamide (Thermo) prior to verification by agarose-gel electrophoresis and/or sequencing. Sequences of used primers are found in excel file S1 and detailed protocols can be download from https://doi.org/10.48420/16878859.

### RNP Assembly and magnetofection and validation

We modified, improved and optimized a previously published simplified forward transfection protocol^45^ by delivering High fidelity Cas9 (Alt-R S.p. HiFi Cas9 Nuclease V3, IDT) and guideRNA (crRNA:tracrRNA) as RNP together (final 10 nM) with ssDNA donor (100-300b homologous arms) via CrisperMAX (Thermo) combined with nanoparticles for magnetofection. Combimag nanoparticles (OZBiosciences) were used according to the manufacturer suggestions, with a magnetic plate. To increase HDR efficiency of shorter homologous arms used for Rab11b knock in (100-150 bases), RAD51-stimulatory compound 1 (RS-1,^46^ was successfully implemented in our protocol (20 uM prior magnetofection), but did not improve efficiency with longer arms used for Rab11a and Rab25 (300 bases). GuideRNA (crRNA:tracrRNA) cutting in a 10bp proximity to the ATG insertion site were selected with a preference for low off target potential. Sequences of tracrRNA and donor plasmids used for ssDNA preparation, all designed to be resistant to Cas9 cleavage, are provided in excel file S1. The amount of long ssDNA (15-105ng) was critical to optimize for each ssDNA preparation method as excess of highly pure full-length ssDNA led to profound cell death and was sometimes accompanied by non-specific knock in events. Knock-in was verified by FP expression and localisation, and FACS sorted cells validated for specific full-length knock in by western blot and selected individual clones were validated by sequencing of PCR-amplified genomic DNA. Primers binding outside the homologous arms used for knock in were used together with mNeonGreen or mCherry specific primers followed by nested PCR with additional primers prior sequencing (sequences of primers are listed in excel file S1). Detailed protocol including screening can be download from https://doi.org/10.48420/16878859.

### Imaging, Colocalization

Low resolution widefield images were acquired using an Olympus IX51/TH4-200 optical microscope with UPLANFL N 4X /0.13 PHL or 20x/0.40 LCAch Infinity-Corrected PhC Phase Contrast Objective; mercury lamp (U-RFL-T power supply). The images were collected using a QImaging® Retiga-SRV CCD digital camera and QCapture software. For live cell imaging, cells were grown in glass bottom culture dishes with #1.5 high performance cover glass coated with FN (10 μg/ml) or in μ-Plate 96 Well Black (#1.5 polymer, tissue culture treated, Ibidi Cat. #89626) and cells imaged in 1x Opti-Klear medium (Marker Gene Technologies Inc) supplemented with 10% (v/v) FCS at 37 °C. Fluorescence high-resolution timelapse images were acquired using a 3i Marianis system with CSU-X1 spinning disc confocal (Yokagowa) on a Zeiss Axio-Observer Z1 microscope with a 63x/1.40 Plan-Apochromat objective, Evolve EMCCD camera (Photometrics) and motorized XYZ stage (ASI). The 405, 488, 561 and 633nm lasers were controlled using an AOTF through the laserstack (Intelligent Imaging Innovations (3I)) allowing both rapid ‘shuttering’ of the laser and attenuation of the laser power. Images were captured using SlideBook 6.0 software (3i). When acquiring 3D optical stacks the confocal software was used to determine the optimal number of Z sections. Maximum intensity projections of these 3D stacks are shown in the results. For Colocalisation analysis of Rab11a and Rab11b, the Colocalization finder tool in ImageJ was used to generate Pearson’s correlation coefficients. Plot profiles were generated by „plot profile“ function in ImageJ and visualized in Prism software. Kymographs were created from time-lapse images using KYmoToolBox in ImageJ. Four colours were visualized using BIOP Lookup Tables (Spring green, Amber, Bright pink, Azure).

A Zeiss LSM 880 Airyscan confocal microscope was used for live imaging triple knock in A2780 cells (mNeonGreen-Rab11a; mCherry-Rab11b; mTagBFP2-Rab25 DEXON ± Alexa-647 labelled Transferrin) using a 63x / 1.46 Plan-Apochromat objective and 1x confocal zoom with pinhole 1 airy unit. Individual emission spectra (mtagBFP2; mNeonGreen, mCherry, Alexa-647) were first determined with a 34 channel spectral detector and 405/488/594/633 lasers, adjusted and collected using an Airyscan detector to prevent any crosstalk [Scanning mode LineSequential; 1024 × 1024]. These images were then loaded to Comdet plugin to simultaneously track all mNeonGreen/ mCherry / mTabBFP2/ Alexa-647 positive vesicles (constant particle size: 6.0; Max distance between colocalized spots: 9.0; 100 cells from 3 independ. exp.). These results were then re-calculated with respect to each channel using a custom-made Python script (provided in 10.6084/m9.figshare.16810546) to map all unique colocalization outcomes and manually visualized as a Venn diagram with the percentage of colocalization for every channel as 100 % total. Images of triple knock in cells recycling Alexa-647 labelled Transferrin, which were spread overnight on 5mm diameter fibronectin-coated coverslips (live or fixed), were collected using a 3i Lattice Light Sheet microscope using a 25x/1.1 water dipping imaging objective with a combination of three diodes: 405/488/560/642nm Bessel beam array (with 100% laser power and Bessel beam length of 50uM and a ORCA Flash V4 CMOS camera (Hamamatsu) with 50ms or 200 ms exposure time for mNeonGreen/mCherry/Alexa-647 or mTagBFP2, respectively. The system was recalibrated for the refractive index of the 1x Opti-Klear medium (Marker Gene Technologies Inc) supplemented with 10% (v/v) FCS before imaging.

LUXon cells were blue-light irradiated before imaging in CO2 incubator with LED Flashlight torch at 25cm distance with high or low mode ((high = 2420 lux (227 FC), low = 613 lux (56 FC)) for the specified time. siR Tubulin (Cytoskeleton, Cat# CY-SC002) was added 30 min before imaging (400 nM). For Alexa647-Transferrin labelling, cells were incubated 10 min on ice, washed with cold Opti-mem media, followed by 30 min incubation at 37 °C with 25 ug/ml Alexa647-Transferrin, another Opti-mem wash, and a change to Opti-Klear™ Live Cell Imaging Buffer for immediate imaging. Hoechst (#3342 Thermo) was also added (5 μg/ml) to Opti-Klear™ media, at least 1h before imaging (no washing).

### Knocksideways

DExCon modified mCherry-Nanobody-Rab11a/b/25 A2780 cells were pre-induced with dox for 72h and magnetofected with GFP-FKBP and pMito-iRFP670-FRB 24 h prior to imaging directly in μ-Plate 96 Well Black (Ibidi Cat. #89626). Cells were imaged in 1x Opti-Klear medium (Marker Gene Technologies Inc) supplemented with Doxycycline (250 ng/ml) and 10% (v/v) FCS with the 3i Marianis CSU-X1 spinning disc system. Images were first taken for randomly chosen cells expressing constructs across multiple imaging areas before rapamycin (200 nM, Sigma) addition. Images were captured at 20 second intervals for approximately 45 min, with the time between rapamycin addition and time-lapse initiation recorded. Timelapses were analysed via ImageJ using custom-made macro automation (the code is provided in https://doi.org/10.6084/m9.figshare.17085632) of the Colocalisation Finder plugin to evaluate the Pearson’s correlation coefficient between mCherry and mitochondrial (pMito) signal over defined areas containing transformed cells using a consistent proportional threshold; this data was expressed both as a series of colocalization graphs plotting the relative intensities of each channel for each pixel as well as a numerical Pearson’s correlation coefficient value. The individual datasets per timelapse were processed by custom outlier detection (provided in https://doi.org/10.6084/m9.figshare.17085632) utilising a gaussian analysis of differences between data points to split the data into subsets of congruent data points, which were then reconnected via subset congruency to produce the single prevailing set of data without outliers. Individual timelapses were scaled per their range to normalise differences between datasets for comparison. This data was then processed as whole data aggregates and individual cells used to generate one-phase association curve fits, minimum and maximum set at 0 and 1 respectively, using Prism software. Calculated halftimes of Rab11 family members generated through individual curve fits were tested for significant differences via Oneway ANOVA with Tukey’s multiple comparisons.

### High throughput imaging and expression/degradation kinetics

Incucyte S3 Live Cell Analysis system, situated in temperature and CO2 controlled incubator, was used for high throughput imaging. Images of cells in phenol red-free RPMI media (Thermo, #11835030) supplemented with 10% v/v foetal bovine serum (FBS) and ciprofloxacin (0.01 mg/ml; Sigma) were acquired with a 10x/1.22 Plan Fluor (wound healing, proliferation) or 20x/ 0.61 S Plan Fluor (fluorescence kinetics) objective and the green and red filter set for fluorescence images with standard exposure settings. Wound healing: Confluent A2780 cells seeded on 96-well ImageLock™ microplates were pre-treated for 24 ± dox (250 ng/ml) ± IAA (100 μg/ml), scratched by the WoundMaker™ to create homogeneous 700-800μm wide wounds and automatically imaged every 1 h in RPMI phenol free media ± dox/IAA using Incucyte® S3 system. This system automatically detected and quantified the ratio of the cell-occupied area to the total area of the initial scratched region (phase-contrast). This data was then processed as whole data aggregates and individual wells (n=6-9 across 3 replicates) were fitted by one-phase association function (Prism software) to calculate the time needed to close 50 % of the normalized wound area. Cell proliferation was analysed on Costar 96 flat bottom plates (Corning™ 3596) by Incucyte software trained to recognize cell occupied area (=confluence %) based on the brightfield images taken every 30 min and its change over 72 hours. At time 0, cells were seeded sparsely (5-10 thousand cells/well) ± dox (250 ng/ml) ± IAA (100 μg/ml) and the logarithm of 2 calculated from the cell confluence normalized to 5 h (starting point of potential dox/IAA effect). Curves slopes were then obtained by linear regression from the linear part of the individual curves (5-40 h; n=6-9 across 3 replicates) and used for calculation of the doubling time [T] by the formula T = log(2)/slope. For fluorescence expression/degradation kinetics analysed by Incucyte, approximately 20 000 cells were seeded per well of Costar 96 flat bottom plate (Corning™ 3596) ± dox/IAA and immediately imaged (red/green/bright) every 30 min (first 2h every 15 min) up to 72 h. Fluorescence intensity was automatically analysed as integrated fluorescence intensity per image divided by phase area confluence change to count for the effect of cell proliferation. To calculate halftime of degradation (expression) kinetics induced by IAA treatment (Dox), data were fitted by one-phase association function. Where appropriate, individual values were scaled per their range as to normalise differences between datasets for comparison (minimum and maximum set at 0 and 1000 respectively).

A BioTek Synergy H1 microplate reader was used to analyze expression kinetics or protein stability of dox-preinduced cells seeded confluent (100 % monolayer) in Costar 96 flat bottom plate (Corning™ 3596) ± dox with Leibovitz’s L-15 phenol red-free (CO_2_ independent media; #21083027 Thermofisher) supplemented with 10% v/v foetal bovine serum (FBS) and ciprofloxacin (0.01 mg/ml). Readings were captured every 1h over 24-48h at 37 °C (gain 100 399/454; gain 100 490/517nm; gain 150 580/610nm). A2780 cells were growing in L-15 media only up to a monolayer and differences in cell numbers were during readings negligible (mTagBFP fluorescence of Teton3G-T2A- mTagBFP). To be able to monitor fluorescence change over 72h, cells were seeded 24h before the experiment ± dox, and fluorescence intensity measured over the following 48h. To account for the phototoxic effect of fluorescence readings and compare differences between datasets, fluorescence kinetics were scaled per their range (minimum and maximum set at 0 and 1000 respectively). Data were aggregated and fitted by one-phase association function (to calculate halftime) or by polynomial function.

Statistics were calculated from halftimes generated through individual curve fits (expression kinetics) or slopes derived from the linear part of the curve (protein stability), unless otherwise indicated.

### Cell-derived matrix and 3D invasion assays

Cell derived matrices were generated as previously described ^29,47^. Briefly, plates were coated with 0.2% gelatin (v/v, Sigma Aldrich), crosslinked with 1% glutaraldehyde (v/v, Sigma Aldrich) and quenched with 1M glycine (Thermo Fisher) before TIFs were confluently seeded. DMEM medium supplemented with 25 μg/ml ascorbic acid (v/v, Sigma Aldrich) was changed every 48h for 8 days. Cells were denuded with extraction buffer (0.5% (v/v) Triton X-100; 20 mM ammonium hydroxide (NH4OH)) to leave only matrix. Finally, phosphodiester linkages in the DNA backbone were cleaved by DNAse I (Lonza).

Collagen/FN matrix for invasion assays was prepared as follows: Collagen solution (10 mg/ml, Corning #354249) was diluted to the final mix: 2.25 mg/ml, 1× RPMI, 15 mM HEPES (750 mM), 8.5 mM NaOH (1 M), 0.4 % NaHCO3 (7.5 %, Sigma), 5 μg/ml folic acid, and supplemented with 25 μg /ml Fibronectin (labelled or un-labelled as needed). FN was dialysed into PBS by Slide-A-Lyzer Dialysis Casette (Thermo) and conjugated with Alexa-fluor 647 NHS ester (Thermo). For LUXon invasion experiments Alexa-647 conjugated FN was then supplemented in a 1:10 ratio (labelled/non-labelled, total 25 μg/ml). For the modified 3D cell-zone exclusion assay ^23^, 50 μl of the collagen/FN (un-labelled) mix was added into well of a 96-well plate and polymerized at 37 °C. Cell suspension (100 μl; 1 × 10^6^ cells/ml) was added on top of the collagen/FN gel. After cell attachment (4 h), the medium was removed and scratch was performed. Immediately after removal of the rest of media in the generated wound, another layer of collagen/FN mix (100 μl, including FN-647) was added (schematic of the experiment is shown in 3I). Collagen was allowed to polymerize for 30 min at RT/37 °C, RPMI medium added and cells incubated for 24h at 37°C in a humidified 5% (v/v) CO2 atmosphere to allow invasion. For spatiotemporal reactivation of Rab25 expression, wells were partly illuminated by blue LED for first 9 hours (black plasticine was used to block light on 80% of the well area).

Spheroids of uniform size were prepared using MicroTissues® 3D Petri Dish® micro-mold spheroids (microtissues; #12-81) for on-chip spheroid invasion assays. Briefly: 600 μl of 2 % UltraPure Agarose (v/v in PBS; Sigma) was allowed to form gels in 3D petri dish micro-mold with 81 circular recesses (9 × 9 array; 800 μm each in diameter). 40 500 cells/190 μl (500 cells/spheroid with an estimated diameter 100 μm) were introduced to the micro-mold and allowed to self-assemble into spheroid over 48hr. To increase the compactness of the spheroid, 10 % serum was changed to 1 % on the second day of spheroid formation (±dox/IAA). Medium was fully aspirated and 190 μl of collage/FN introduced and polymerised at 37C. Chip with 81 spheroids embedded in collagen/FN gel were overlayed with 0.8-1% UltraPure Agarose (v/v in PBS) to prevent gel movement and shrinking. Finally, chips were overlayed with RPMI media (10 % serum; ±dox/IAA 2x excess to count for the volume of agarose), imaged (4x objective, phase contrast) and cells allowed to invade for 48-72 h at 37°C in a humidified 5% (v/v) CO2 atmosphere (±blue light illumination) before additional imaging. Detailed protocol can be download from https://doi.org/10.48420/16878859. Cell invasion area was calculated custom-written macro code for ImageJ that automatically created a binary mask of invading cells and subtracted the initial spheroid area (provided in https://doi.org/10.48420/16878829).

### Statistical analysis

Data were tested for normality and one-way ANOVA with Tukey post hoc test used for multiple comparisons as indicated in legend. Where data were not normally distributed, ANOVA on ranks was used instead. All statistical analysis has been performed with GraphPad Prism software, where *** denotes p < 0.001, ** denotes p < 0.01, and * denotes p < 0.05. Data represent at least three independent experiments; n numbers and p values are described in relevant figure legends.

### Data availability

List of antibodies, chemicals, materials, primers and all necessary sequences for replication of this study are provided (excel file S1 or accessible from https://doi.org/10.48420/14999550), along Graph Prism files (raw or normalized data at https://doi.org/10.48420/16878904), detailed innovative protocols 10.48420/16878859 and custom scripts/macro codes are deposited on Figshare (links in the methods section). Annotated plasmid maps used and generated in this study (including catalogue of donors) can be download from 10.48420/16810525 and 10.48420/14999526.

Plasmids can be provided upon reasonable request from corresponding author or will be available from Addgene. Supplementary movies can be downloaded from https://doi.org/10.48420/17111864.

## Acknowledgement

We thank Hayley Bennett for providing a protocol for long ssDNA preparation using biotinylated primers and all of the members of the Caswell lab for their support, especially Eleanor Hinde. We also are grateful to Itaru Imayoshi and Mayumi Yamada for providing lentiviral plasmids coding two versions of PA-Tet-OFF/ON. The Bioimaging Facility microscopes used in this study were purchased with grants from BBSRC, Wellcome and the University of Manchester Strategic Fund. We thank **Peter March**, **Steven Marsden** and Dave Spiller for their help with microscopy. We further thank the University of Manchester Flow Cytometry Core Facility for assistance with flow cytometry and sorting. The Flow Cytometry Core Facility is supported in part by the University of Manchester with assistance from MRC Grant ref MR/L011840/1. This project has received funding from the European Union’s Horizon 2020 research and innovation programme under grant agreement No [836212]. This work was further supported by the MRC (MR/R009376/1), CRUK (DCRPGF\100002) and the Wellcome Trust Centre for Cell Matrix research is funded by grant 203128/A/16/Z. GTEx project was supported by the Common Fund of the Office of the Director of the National Institutes of Health, and by NCI, NHGRI, NHLBI, NIDA, NIMH, and NINDS.

## Author Contributions

Author contributions: Conception and design of the work, J.G., P.C., A.A.; acquisition, analysis and interpretation of data, J. G., T.H, C.F., P.C; writing – original draft: J.G., P.C; writing – review and editing: all authors.

## Supplementary figures

**Figure S1.**
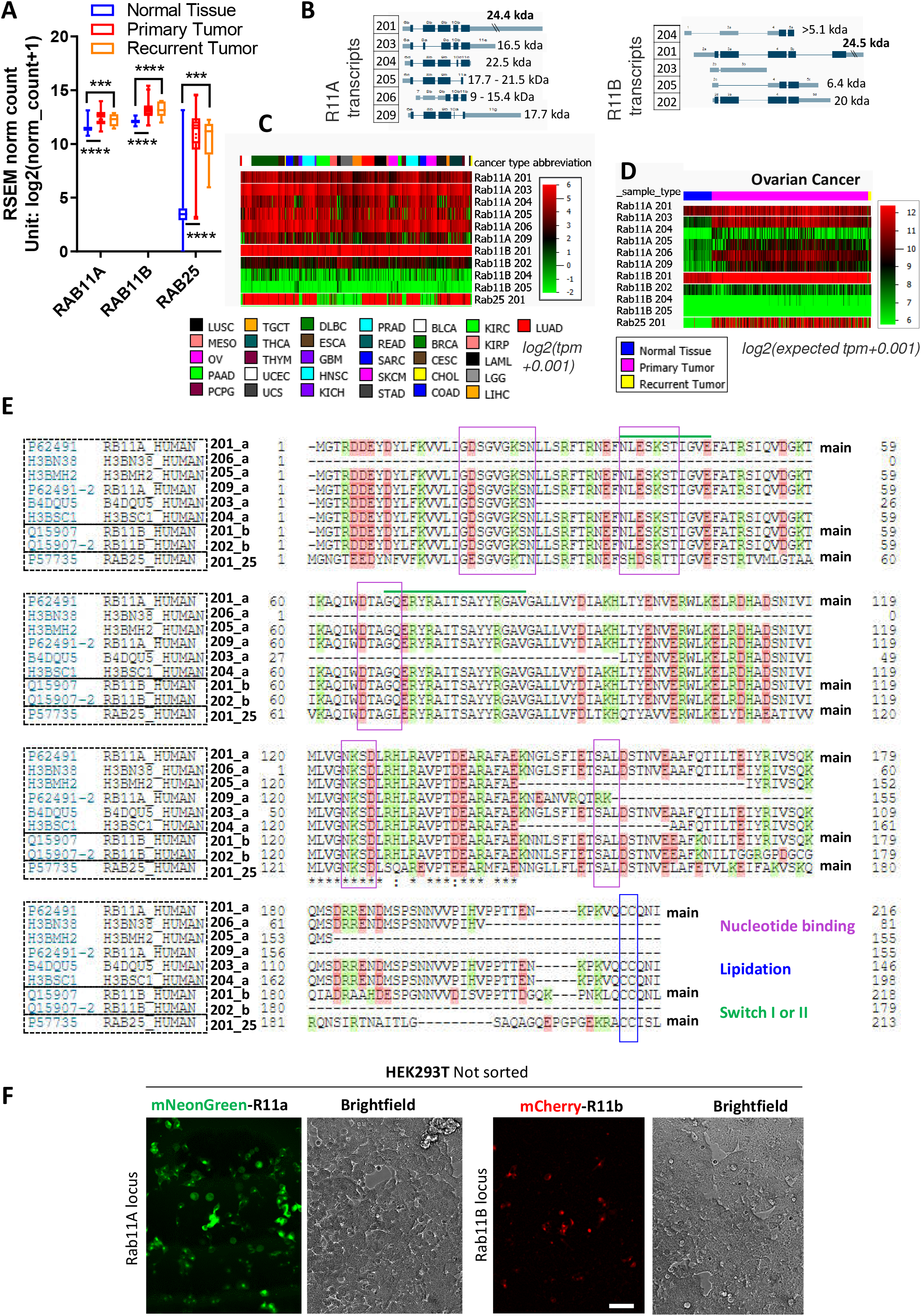
Rab11 family expression profiling. A) Comparison of Rab11s expression in normal healthy or primary and recurrent ovarian tumors (TCGA TARGET GTEx study). Schematic illustration of all predicted alternatively spliced protein coding variants of Rab11a/b transcripts with corresponding introns/exons. C-D) Comparison of Rab11a/b/25 isoform expression levels (RNAseq) across multiple types of cancer (TCGA PanCan dataset) C) and recurrent ovarian tumors from the TCGA TARGET GTEx study D). E) Multiple sequence alignment (made by UNIPROT) of the most abundant protein-coding Rab11s transcripts. Sequences important for GTP/GDP binding or for the attachment to the membrane (CC) are highlighted. D) Fluorescence widefield images of non-sorted mNeonGreen or mCherry knock ins to Rab11Aor Rab11b loci of HEK293T. Scale bar=100 μm.

**Figure S2.**
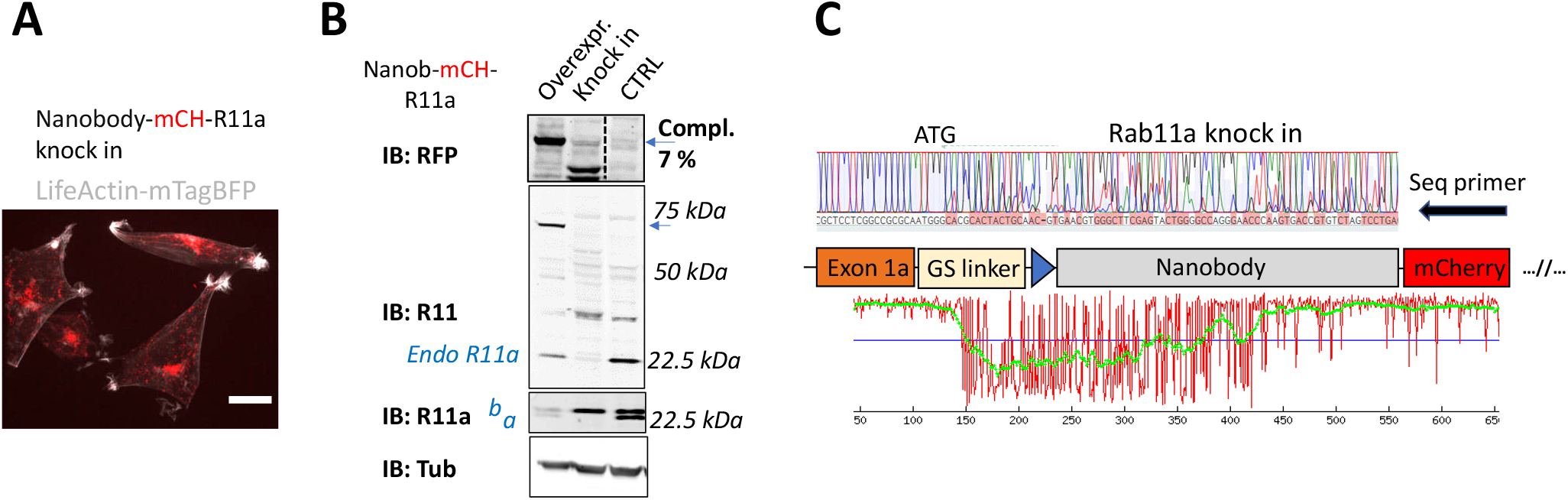
N-terminal knock-in of non-fluorescent coding gene and validation. A) Spinning disk confocal image of A2780 with antiGFPnanobody-mCherry knock in to Rab11a locus stably expressing LifeActin-mtagBFP (grey). Signal of mCherry (red; = mCH) reports full length integration only of mCherry, not of antiGFPnanobody (scale bar=20 μm). B) Immunoblots of A2780 wt (CTRL), antiGFPnanobody-mCherry knock in to Rab11a locus or A2780 overexpressing stably integrated full length antiGFPnanobody-mCherry-Rab11a (lentiviral transduction, expected protein size 69 kDa marked by blue arrow). Blots were probed with anti-mCherry (RFP), anti-Rab11a or anti-Rab11 (targeting both Rab11a/b) antibodies (a = Rab11a; b = Rab11b). Tubulin (Tub), loading control. C) Chromatogram of mCherry-antiGFPnanobody-Rab11a cells (classical knock in) with schematic of knock in outcome (seq. prim = sequencing primer). Graph, made by TIDER analysis, estimates the frequency of mutations/indels in knocked in cells^50^.

**Figure S3.**
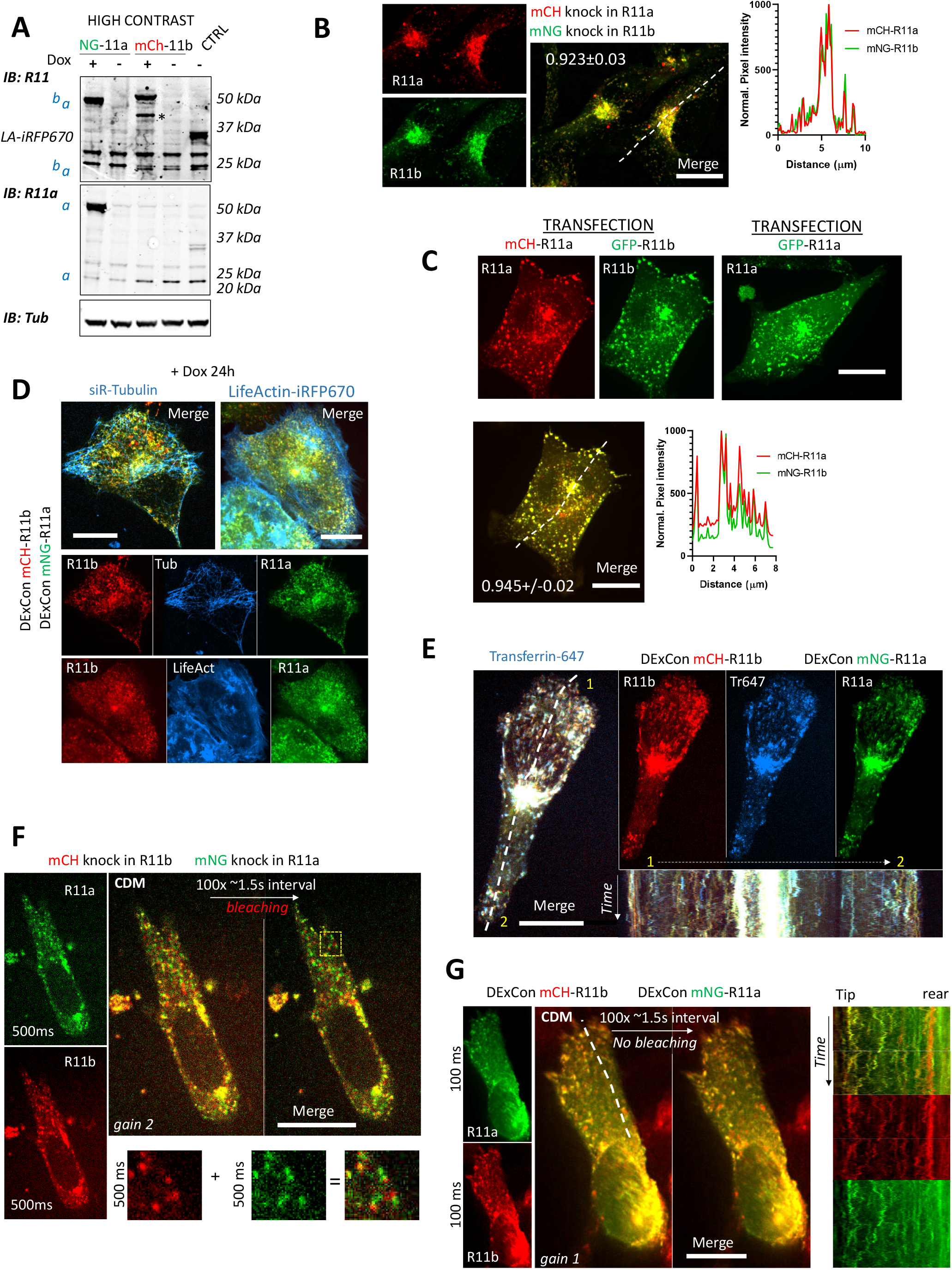
DExCon controls Rab11a and Rab11b expression and preserves physiological localization. A) Higher contrast immunoblots as in 2B. B-G) Spinning disk confocal images; see movie S1-4. B) Classical double mNeonGreen or mCherry knock ins to Rab11a or Rab11b as indicated. C) A2780 co-transfected (magnetofected) with cDNA for mCherry-Rab11a and GFP-Rab11b; line profile corresponds to the yellow dashed line and number reflect Pearson cross correlation coefficient (average ± SEM, n = 24 cells; maximum intensity z-projection). D-E) Double DExCon mNeonGreen-Rab11a / mCherry-Rab11b stably expressing LifeActin-iRFP670 (D, right) or treated with siR-Tubulin (400 nM, (D, left)) or E) following uptake of ALEXA-647 labelled Transferrin (kymograph corresponds to dashed line; total 5 min). F-G) Spinning disk confocal images of cells migrating in <u>C</u>ell <u>D</u>erived <u>M</u>atrix acquired with same settings (only gain and exposure settings different as indicated); see movie S4. F) mNeonGreen-Rab11a / mCherry-Rab11b double knock in A2780 with endogenous promoter. Bottom images (zoom of yellow rectangle) highlight the imaging delay artefact of vesicle de-colocalization. G) Double DExCon mNeonGreen-Rab11a / mCherry-Rab11b imaging of cells. On the right corresponds to the white dashed line; total 4.4 min. B-G) Scale bar=20 μm.

**Figure S4.**
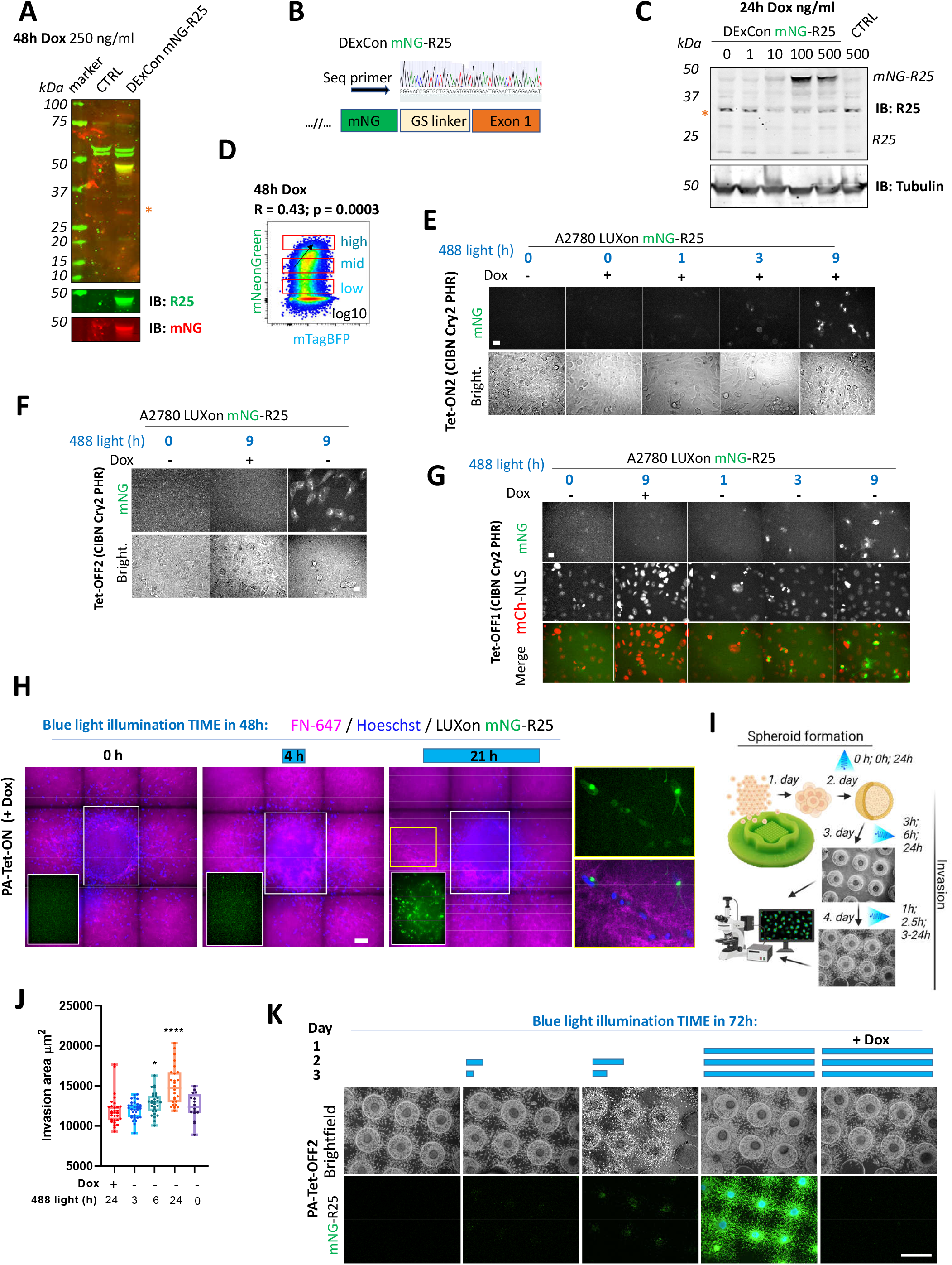
Reactivation of Rab25 expression. A) Immunoblots of A2780 cell lysates (CTRL) or mNeonGreen-Rab25 DExCon A2780 cells treated with dox as indicated (250 ng/ml) probed with anti-Rab25 or anti-mNeonGreen (red stars indicate unspecific band always recognized by secondary antibody). Tubulin, loading control, CTRL un-modified A2780. B) Chromatogram of mNeonGreen-Rab25 DExCon A2780 cells with schematic of knock in outcome (seq. prim = sequencing primer). C) Immunoblots as in A). D) FACS of mNeonGreen-Rab25 DExCon cells re-sorted for different expression levels of Rab25. Black arrow indicates positive correlation of mNeonGreen (Rab25) and mTagBFP (Teton3G) levels; R = Pearson’ cross-correlation coefficient; p-value. E-H) Spinning disc confocal images of mNeonGreen-Rab25 LUXon cells (Rab25 as mNG) expressing PA-Tet-ON or PA-Tet-OFF or their versions with an improved blue light sensitivity (ON2; OFF2) treated by dox and blue light (26.8 W/m2) as indicated. scale bar=20 μm. In case of G) mCherry-NLS (nuclear reporter) was used. H) Spheroids are labelled with Hoechst 3342 and allowed to invade (48 h) collagen matrix supplemented with labelled FN-647. Three conditions of blue light illumination (1^st^+2^nd^ day) are shown as merge of all three channels, white rectangle for the mNeonGreen channel only, or zoom of the yellow rectangle. I) Schematic of spheroid “on chip” invasion assay with illumination protocol used in J-K) with mNeonGreen-Rab25 LUXon (PA-Tet-OFF2) cells. J) Quantification (n = 16-34 per each condition from 3 independent experiments, one way ANOVA Tukey post hoc test), K) Representative fluorescence images (scale bar=1 mm). Schematic illustration was created with BioRender.com.

**Figure S5.**
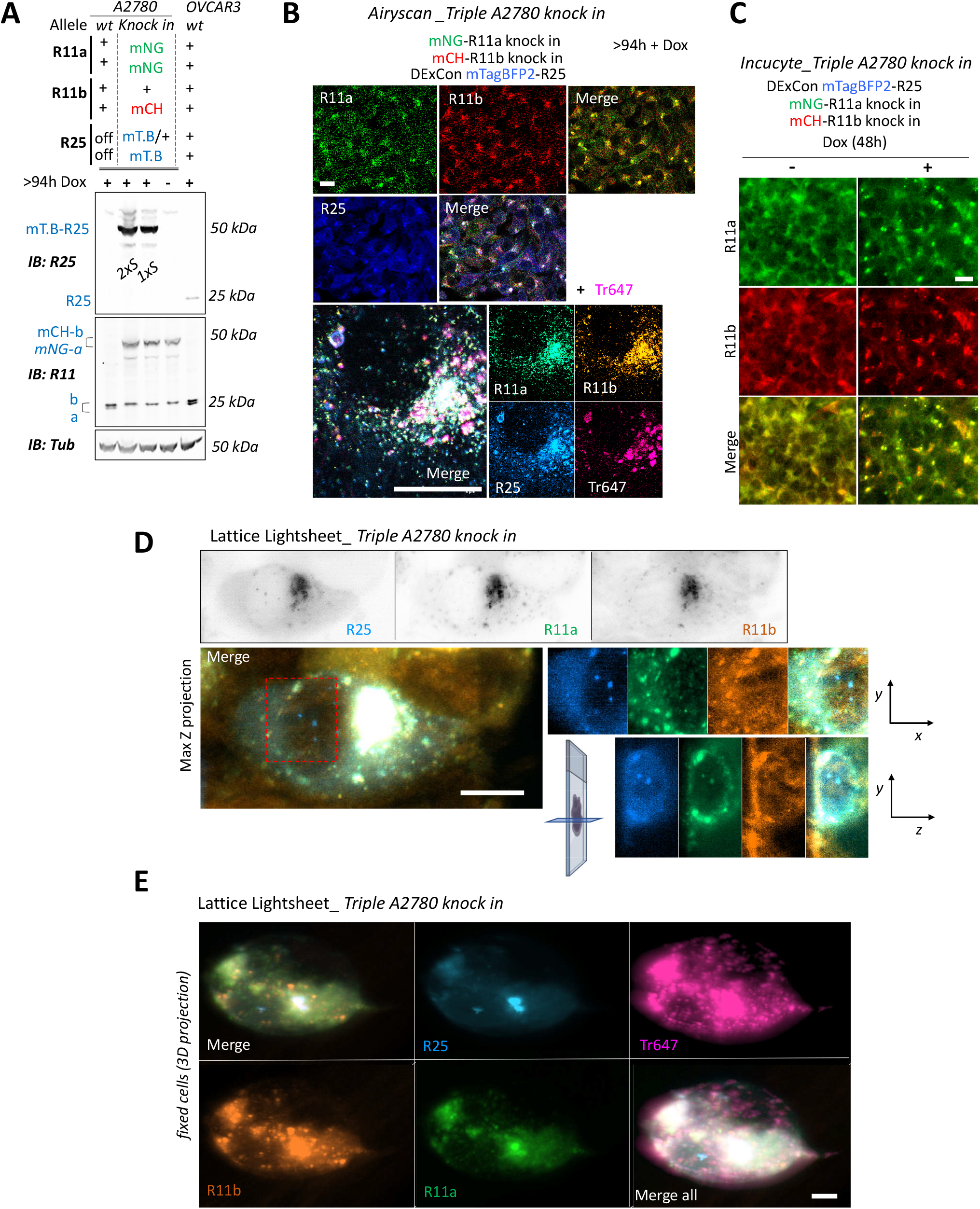
mNeonGreen-Rab11a, mCherry-Rab11b, DExCon-mTagBFP2-Rab25 triple knock-in. A-E) Triple knock in A2780 cells (mNeonGreen-Rab11a; mCherry-Rab11b; mTagBFP2-Rab25 DExCon) treated by dox (>94 h); B-E) recycling Alexa-647 labelled Transferrin (Tr647; 30 minutes). A) Immunoblots. Fluorescent antibodies: anti-Rab11 targeting both Rab11a/b (Rab11), anti-Rab25 (R25) shown as black and white; Tubulin (Tub), loading control. wt= un-modified A2780 or OVCAR3; a = Rab11a; b = Rab11b (heterozygote); R25 = Rab25 ± fluorophore knock in. Sorted (1 or 2xS). B) AiryScan fluorescence images (LSM880; 63x; Scale bar 20 μm). C) Images obtained by Incucyte® S3 system (Scale bar 20 μm). D) Lattice LightSheet images (3i; Scale bar 10 μm; live cells); one focal plane or Maximum Intensity Z-projection for merged channels with increased contrast. See also 4F and movie S11. E) Lattice LightSheet images (3i; Scale bar 5 μm) of fixed cells (3D projection using Imaris Cell Imaging Software); See also movie S12.

**Figure S6.**
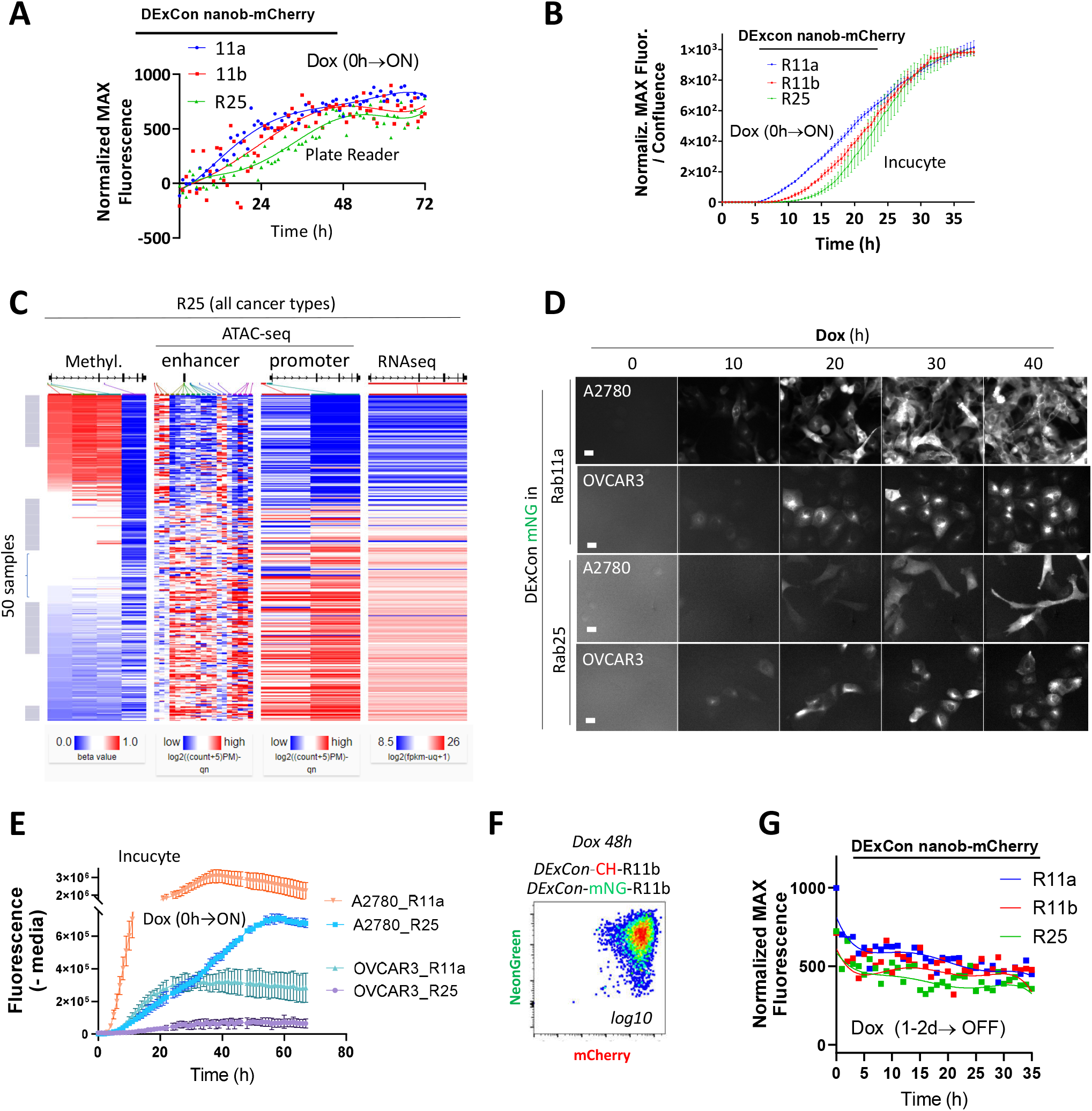
Protein expression kinetics and stability are revealed using DExCon. A-B) Comparison of expression kinetics of DExCon antiGFPnanobody-mCherry-Rab11s by microplate reader (A) or Incucyte® S3 (B). Representative examples are shown from 3 independent experiments; A) fitted by fifth order polynomial function; cells seeded confluent fluorescence normalized to the maximal intensity. B) Mean ± SEM; normalized to maximal fluorescence intensity and cell confluence. C) Bioinformatic analysis of Rab25 in all cancer types using UCSC Xena; GDC Pan-Cancer (PANCAN) study: DNA promoter methylation (Methylation450K), RNA expression (RNAseq) and compactness of chromatin structure in its promoter and enhancer regions (ATAC-seq). D) Representative fluorescence timelapse images of DExCon mNeonGreen-Rab11a or -Rab25, generated in A2780 or OVCAR-3 cells, following dox induction (Incucyte® S3) and E) absolute fluorescence intensity quantification normalized to cell confluence (Incucyte® S3). See also 5D. F) FACS of mNeonGreen/mCherry Rab11b double DExCon cells (dox 48h; 2x sorted). See also 5G. G) Comparison of DExCon antiGFPnanobody-mCherry-Rab11s (A2780) protein stability. Cells pre-induced by dox for 24-48h followed by dox removal and fluorescence determined by microplate reader. Cells were seeded confluent and fluorescence normalized to the maximal intensity (1000). Curves fitted by fifth order polynomial function are shown (n=12; 4 independent experiments).

**Figure S7.**
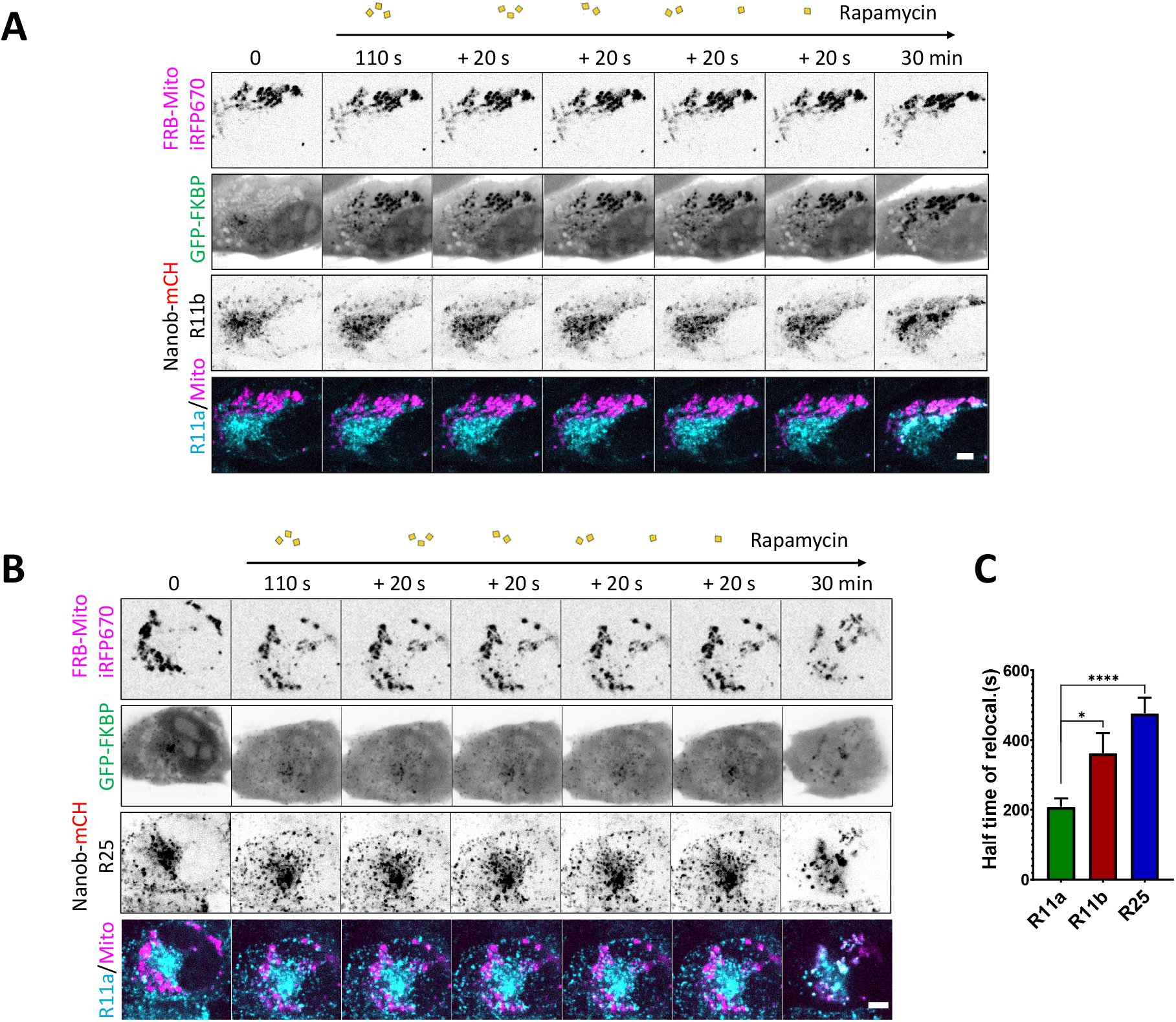
Protein dynamics are revealed using DExCon. A-B) Representative confocal fluorescence timelapse images of antiGFPnanobody-mCherry-R11b A) or –Rab25 B) DExCon cells induced with dox for 72 h (see also 5K, L). Cells were co-magnetofected with FRB-Mito-iRFP670 and GFP-FKBP on 96 well ibidi imaging plate 24 hours before imaging and their heterodimerization induced by 200 nM Rapamycin as indicated. Scale bar=5μm. C). Halftime of Rab11a/b/25 relocalisation were fitted (see 5M) and quantified. Bars represent mean ± SEM; n = 10-18 from 3-4 independent experiments, one way ANOVA with Tukey post hoc test used for statistical analysis.

**Figure S8.**
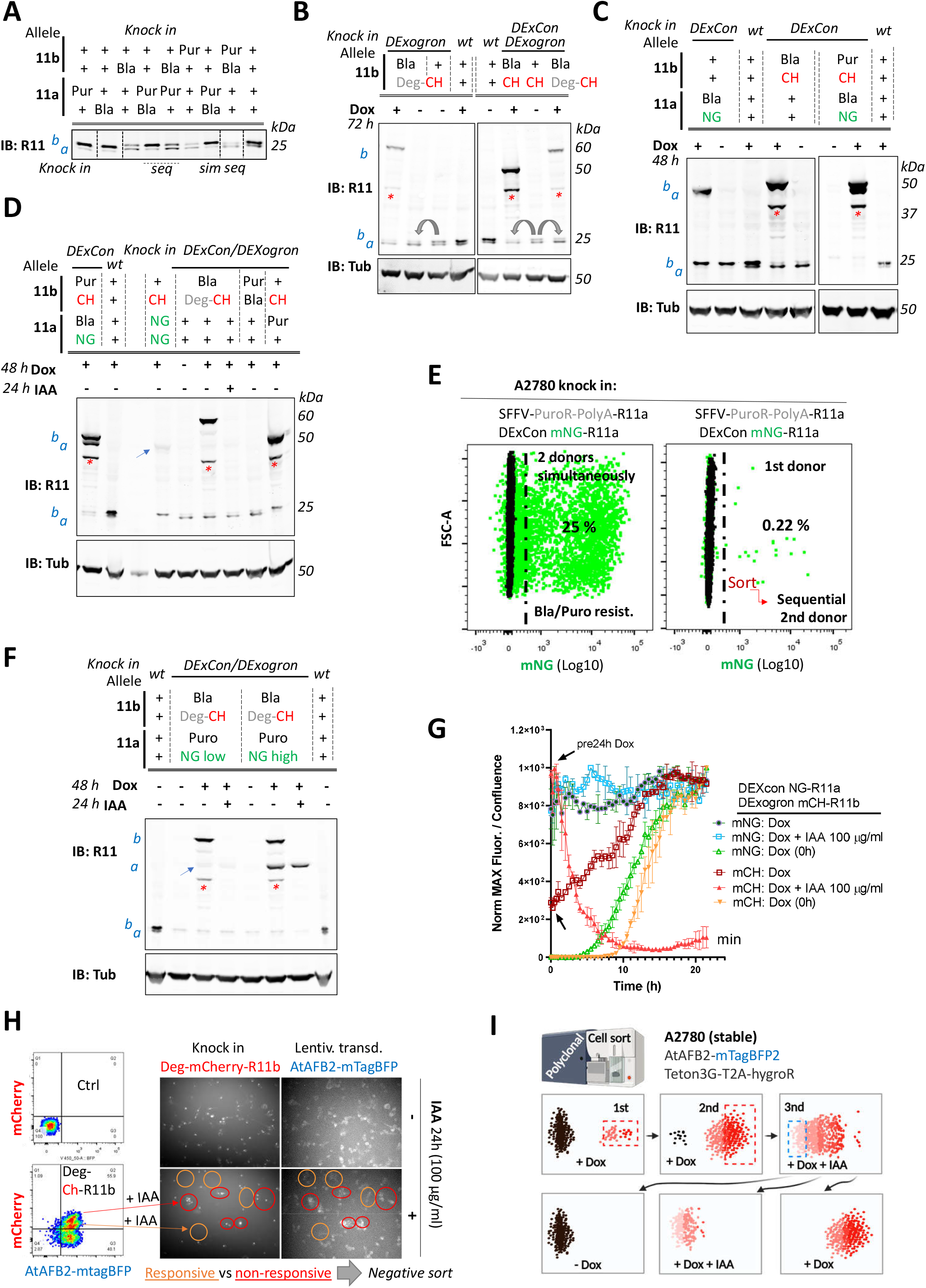
The DExogron module can tune protein levels within cells. A-D, F) A2780 cells were CRISPR modified to introduce different cassettes to Rab11a, Rab11b or both loci. Different alleles were targeted as indicated (± DExCon); Puro/Bla: Puromycin or Blasticidine resistance were introduced with SFFV promoter and PolyA (see 6B) and led to DExCon or DExogron knock in improvement (grey arrows). Immunoblots were probed with anti-Rab11 antibody targeting both Rab11a/b (Rab11). Tubulin (Tub), loading control. wt= un-modified A2780; a = Rab11a; b = Rab11b ±FP knock in (blue arrows indicate mNeonGreen-Rab11a DExCon band sorted for low or endogenously tagged mNeonGreen/mCHerry-Rab11a or b. Red stars indicate mCherry/iRFP670 lower molecular weight band caused by hydrolysis during sample preparation^20^. miniIAA7 = Deg; CH = mCherry; NG = mNeonGreen. Cells were treated ±dox ±IAA as indicated. Sequential (seq) or simultaneous (sim) knock in of two different donors is highlighted in A) and compared for percentage of cells expressing mNeonGreen-Rab11a DExCon by FACS E). Percentage of mNeonGreen positive cells (left selected for puromycin resistance) is shown after CRISPR based knock-in in green. Black: unmodified A2780 Ctrl. G) Expression and degradation kinetics of Dexogron-Cherry-Rab11b/DExCon-mNeonGreen-Rab11a cells (A2780) analysed by Incucyte imaging. Cells treated ±dox (250 ng/ml) ± IAA as indicated (arrows indicate dox stimulation, 24h before analysis). Curves normalized to maximal fluorescence intensity and cell confluence, mean of 3 independent experiments ± SD is shown. H) FACS and fluorescence images of miniIAA7-mCherry-Rab11b knock-in A2780 cells stably expressing AtAFB2-mTagBFP (2x sorted) ± IAA as indicated. IAA responsive (middle population, left bottom, orange ovals) and non-responsive cells (top population, left bottom, red ovals) are indicated. Ctrl = un-modified A2780. I) Pipeline that maintains polyclonal CRISPR modified cells (three cycles of sort, first two for the knock-in cell enrichment followed by negative sort to obtained IAA responsive cells). Illustration created with BioRender.com.

**Figure S9.**
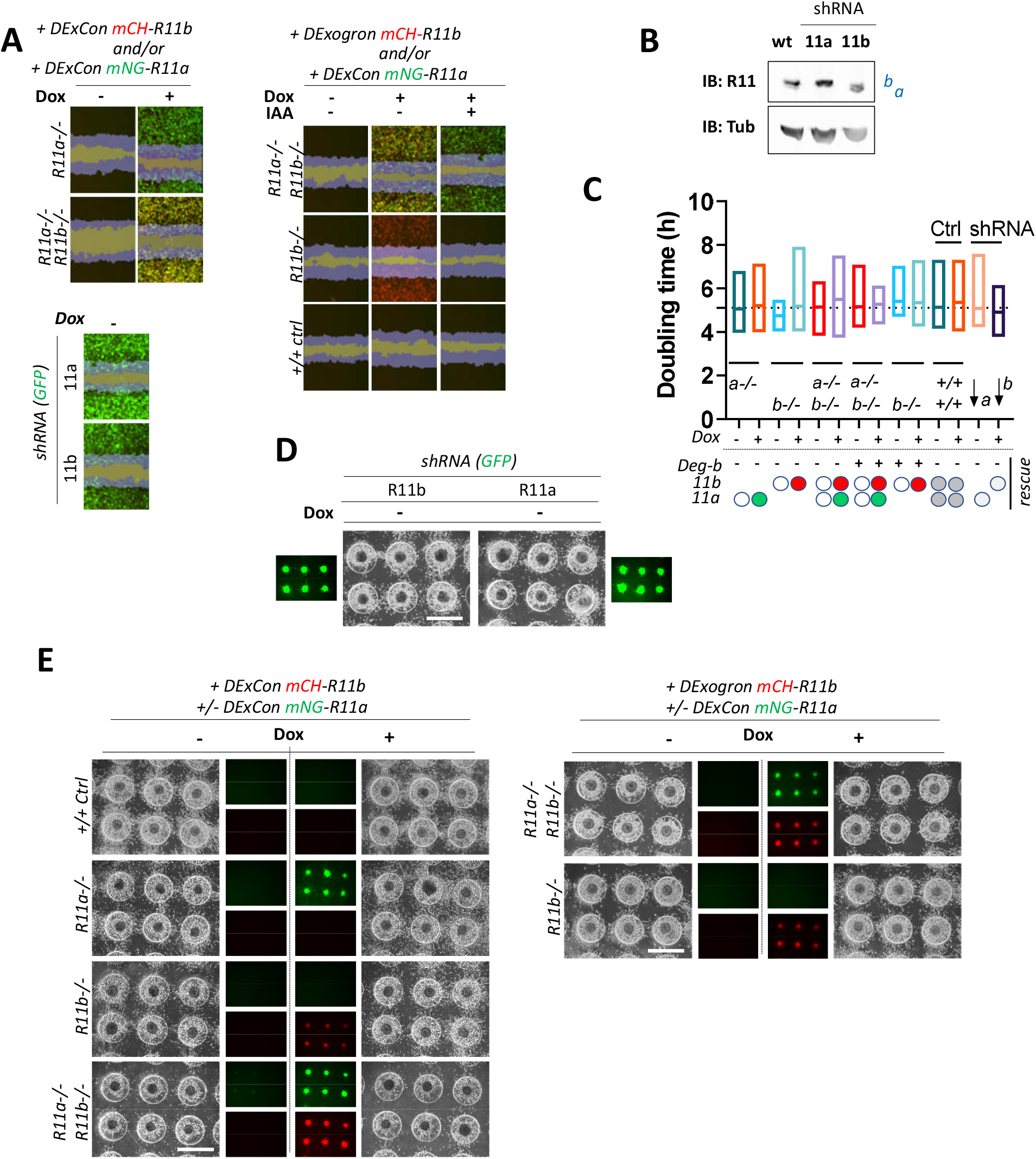
DExCon/DExogron reveal the contribution of Rab11a/b to migration and invasion. A-E) a-/-:DExCon-mNeonGreen-Rab11a (rescue shown as green oval or as green fluorescence), b-/-: DExCon-mCherry-Rab11b (rescue shown as red oval or as red fluorescence) or Deg-b: DExogron-mCherry-Rab11b; Ctrl +/+ as A2780 targeted by shRNA anti-Rab11a or b (see arrows). A) Confluent cells were pre-treated for 24 ± dox (250 ng/ml) ± IAA (100 μg/ml), scratch wounds introduced and imaged in phenol free RPMI±dox/IAA as indicated. Scratch wound migration experiments were imaged and analysed by Incucyte (blue mask shown based on brightfield image taken at time 0 and 6 h after the scratch together with mCherry=red/mNeonGreen=green; fluorescence; yellow = double rescue). For quantification see 6F). Movie S16. B) Immunoblot of Rab11a/b shRNA in A2780 cells. Blots were probed with anti-Rab11 antibody targeting both Rab11a/b (Rab11). C). Proliferation rate calculated as doubling time (h) from brightfield images generated and automatically analysed as cell confluence area change (0-72 h) using Incucyte. D-E) On-chip spheroid invasion assays. Cells migrated into collagen matrix supplemented with FN and treated ± dox (250 ng/ml) as indicated (24h pre-induced). See also 6H and for quantification 6G, I. Brightfield and fluorescence images (scale bar=1mm).

**Figure S10.**
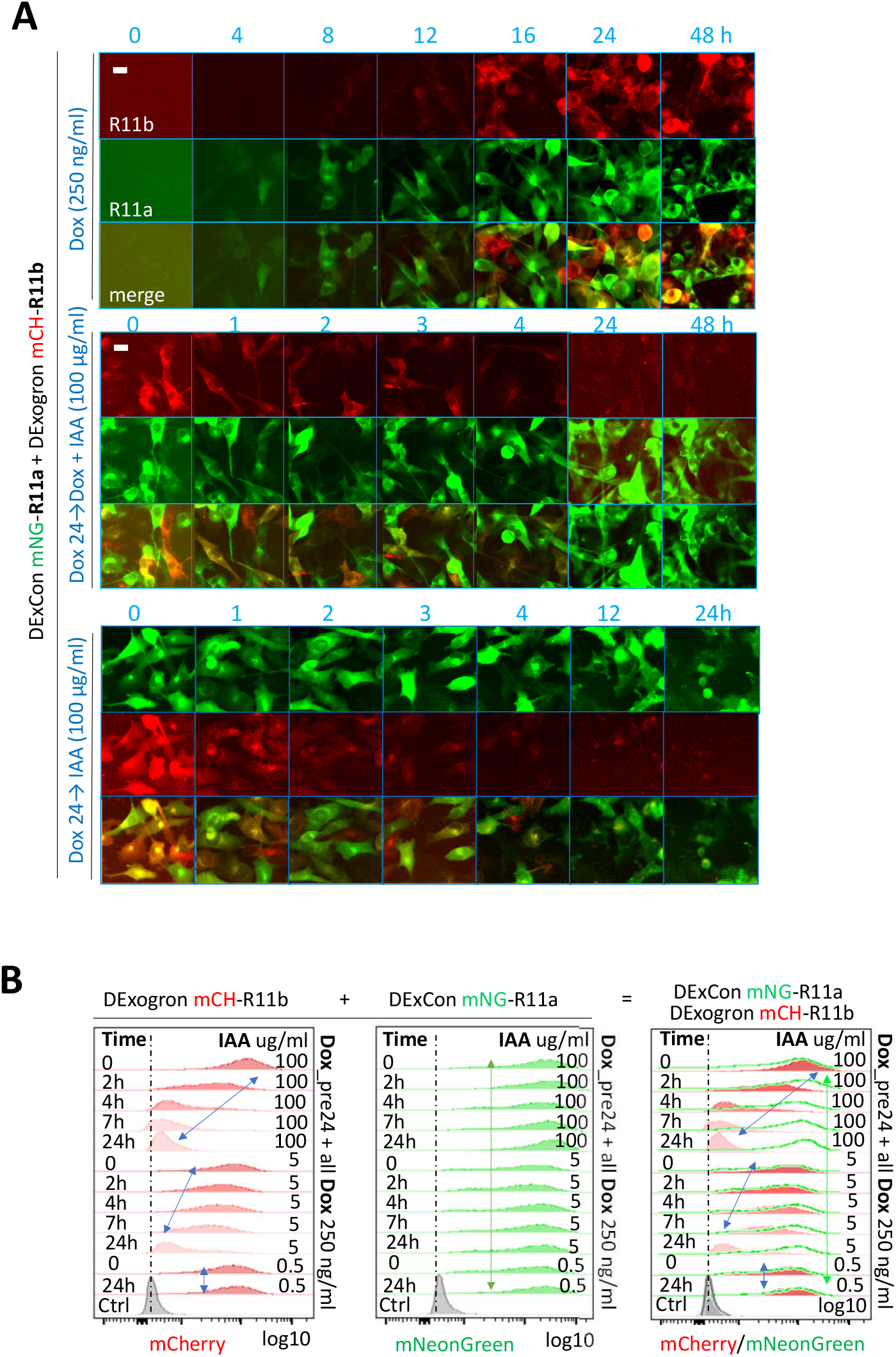
Characterisation of DExCon-Rab11a/DExogron-Rab11b cells. A-B) DExCon-Rab11a/ DExogron-Rab11b cells treated with IAA, dox or dox/IAA as indicated; dox=250 ng/ml, IAA=100 μg/ml unless otherwise stated. Dox (pre)24=dox stimulation 24h before analysis. mCH = mCherry; mNG = mNeonGreen. A) Timelapse fluorescence images obtained automatically by Incucyte (scale bar=20μm). B) FACS analysis, arrows indicate dynamic range and change in fluorescence intensity of mCherry or mNeonGreen; Ctrl = un-modified A2780 highlighted by black (max. peak) and blue dashed lines (gaussian signal distribution).

**Figure S11.**
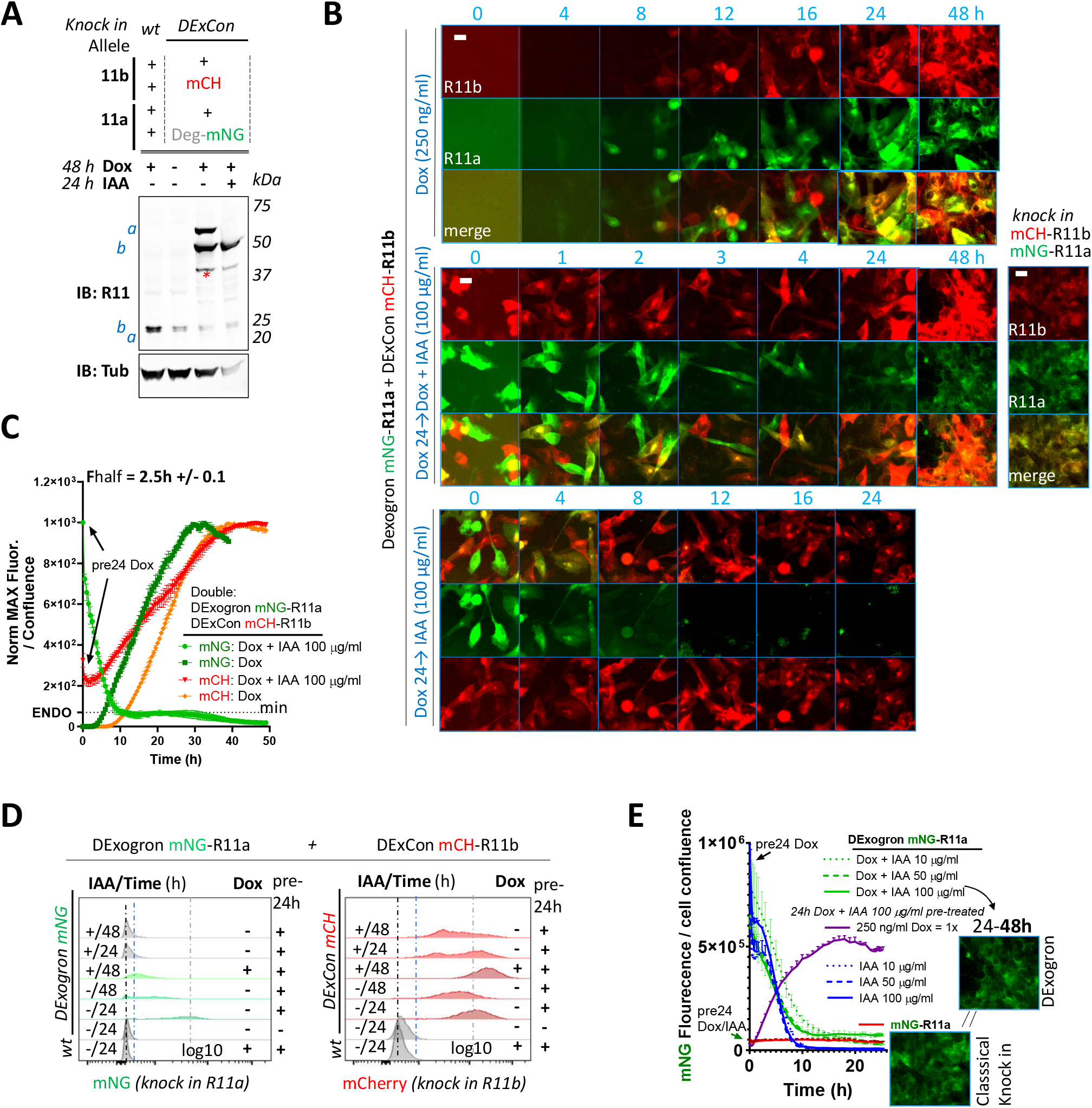
Characterisation of DExogron-Rab11a/DExCon-Rab11b cells. A-E) DExogron-mNeonGreen-Rab11a/DExCon-mCherry-Rab11b A2780 (single clone) cells treated with IAA, dox or dox/IAA as indicated; dox=250 ng/ml, IAA=100 μg/ml unless otherwise stated. Dox (pre)24=dox stimulation 24h before analysis. mCherry-Rab11b/mNeonGreen-R11a conventional knock-ins are shown for comparison (right). mCH = mCherry; mNG = mNeonGreen; miniIAA7 = Deg. A) Immunoblots probed with anti-Rab11 antibody targeting both Rab11a/b (Rab11); Tubulin (Tub), loading control. wt= un-modified A2780; a = Rab11a; b = Rab11b ± fluorophore knock in. Red stars indicate mCherry/iRFP670 lower molecular weight band caused by hydrolysis during sample preparation^20^. B) Timelapse fluorescence images obtained using Incucyte (scale bar=20μm). C) Expression and degradation kinetics analysed by Incucyte. Arrows indicate dox stimulation 24h before analysis. Curves were normalized to maximal fluorescence intensity and cell confluence, mean of 3 independent experiments ±SD shown. Half-time of mNeonGreen fluorescence decrease to the fluorescence intensity of conventional mNeonGreen-Rab11a knock in (dotted line marked as ENDO, see also E) calculated as 1.7h ± 0.1 (mean of 3 independent experiments ±SEM) D). FACS analysis of fluorescence intensity of mCherry or mNeonGreen (grey dashed line for 24h pre-induced reference); Ctrl = un-modified A2780 highlighted by black (max. peak) and blue dashed lines (gaussian signal distribution). E) Expression and degradation kinetics analysed by Incucyte. Curves were normalized to cell confluence, mean of 3 independent experiments ±SEM shown. See movie S17. Images represent conventional mNeonGreen-R11a knock in fluorescence intensity (bottom) and DExogron-mNeonGreen-Rab11a co-stimulated with dox/IAA for 24-48 h (on the right).

## Supplementary movie legends

Included movies (https://doi.org/10.48420/17111864) are separate files relevant to the study:

Movies S1-S4 show the advantage of mNeonGreen-Rab11a / mCherry-Rab11b double DExCon (A2780) cells for fluorescence live-cell imaging microscopy (Spinning Disc 3i). Movies S5-S7 show spatiotemporal control of mNeonGreen-Rab25 LUXon cells (A2780) with silenced Rab25 re-activated from genome using blue light (Spinning Disc 3i). Movies S8-S12 show triple knock in A2780 cells (mNeonGreen-Rab11a; mCherry-Rab11b; mTagBFP2-Rab25 DExCon) treated by dox (>94 h); D-F) recycling Alexa-647 labelled Transferrin (Tr647; 30 minutes). Imaging by AiryScan LSM880 or by 3i Lattice LightSheet microscope. Movies S13-S15 and S17 show protein expression kinetics of Rab11s DExCons and degradation kinetics of Rab11s DExogrons (Incucyte® S3 system). Movie S16 shows wound healing experiment automatically imaged and analysed in real time by Incucyte® S3 system of cells as indicated (Rab11s DExCons / DExogrons; shRNA anti-Rab11a or b; A2780). https://doi.org/10.48420/17111864.

### Supplementary movie S1

Timelapse (Spinning disc 3i, 63x) of mNeonGreen-Rab11a / mCherry-Rab11b double DExCon cells (A2780, dox for 24h) treated by siR-Tubulin (400 nM). Timelapse covers total 5 min with frame taken every 2.67s (approximately 10.3s elapsed time per second of the movie); μ-Plate 96 Well Black (Ibidi cat.No 89626; #1.5 polymer coverslip, tissue culture treated; Opti-Klear™ Live Cell Imaging Buffer). Selected frame from this movie is shown in S3D.

### Supplementary movie S2

Timelapse (Spinning disc 3i, 63x) of mNeonGreen-Rab11a / mCherry-Rab11b double DExCon cells (A2780, dox for 24h) stably expressing LifeActin-iRFP670. Timelapse covers total 7 min with frame taken every 2s (approximately 14.5s elapsed time per second of the movie); μ-Plate 96 Well Black (Ibidi cat.No 89626; #1.5 polymer coverslip, tissue culture treated; Opti-Klear™ Live Cell Imaging Buffer). Selected frame from this movie is shown in S3D.

### Supplementary movie S3

Timelapse (Spinning disc 3i, 63x) of mNeonGreen-Rab11a / mCherry-Rab11b double DExCon cells (A2780, dox for 24h) recycling ALEXA-647 labelled Transferrin (25 ug/ml). Timelapse covers total 5 min with frame taken every 2.67s (approximately 16.2s elapsed time per second of the movie); μ-Plate 96 Well Black (Ibidi cat.No 89626; #1.5 polymer coverslip, tissue culture treated; Opti-Klear™ Live Cell Imaging Buffer). Selected frame from this movie is shown in S3E.

### Supplementary movie S4

Timelapse (Spinning disc 3i, 63x) of mNeonGreen-Rab11a / mCherry-Rab11b double knock in A2780 cells ± TRE3GS promoter (DExCon; ±dox 24h) migrating in Cell Derived Matrix (3D). Timelapse covers total 2.25 min (left) or 4.4 min (right) with frame taken every ~1.5s for both (approximately 10.1s elapsed time per second of the movie); μ-Plate 96 Well Black (Ibidi cat.No 89626; #1.5 polymer coverslip, tissue culture treated; Opti-Klear™ Live Cell Imaging Buffer). Selected frames from this movie are shown in S3F-G.

### Supplementary movie S5

Timelapse (Spinning disc 3i, 63x) of mNeonGreen-Rab25 LUXon cells (Rab25 as green) expressing PA-Tet-OFF with mCherry-NLS reporter (red) 18h after being illuminated by blue light (10 hours). Timelapse covers total 1 min with frame taken every 1s (approximately 6.7s elapsed time per second of the movie); μ-Plate 96 Well Black (Ibidi cat.No 89626; #1.5 polymer coverslip, tissue culture treated; Opti-Klear™ Live Cell Imaging Buffer). Selected frame from this movie is shown in 3H.

### Supplementary movie S6

3D projection (Spinning disc 3i, 63x) of mNeonGreen-Rab25 LUXon cells (Rab25 as green) expressing PA-Tet-OFF with mCherry-NLS reporter (red) 18h after being illuminated by blue light (10 hours). μ-Plate 96 Well Black (Ibidi cat.No 89626; #1.5 polymer coverslip, tissue culture treated). Selected frame from this movie is shown in 3H.

### Supplementary movie S7

3D projection (Spinning disc 3i, 63x) of mNeonGreen-Rab25 LUXon cells (Rab25 as blue) expressing PA-Tet-OFF with mCherry-NLS reporter (red) 20h after being spatiotemporally illuminated by blue light (10 hours). Cells are invading to cell-free collagen matrix labelled by FN-647 (green) while being illuminated by blue light of varying intensity (see experimental set-up, right corner). μ-Plate 96 Well Black (Ibidi cat.No 89626; #1.5 polymer coverslip, tissue culture treated) partly covered by black plasticine. Selected frame from this movie is shown in 3J.

### Supplementary movie S8

3D projection of Triple knock in A2780 cells (mNeonGreen-Rab11a; mCherry-Rab11b; mTagBFP2-Rab25 DExCon) treated by dox (>94 h). μ-Plate 96 Well Black (Ibidi cat.No 89626; #1.5 polymer coverslip, tissue culture treated; Opti-Klear™ Live Cell Imaging Buffer). AiryScan LSM880 (63x). Selected frames from this movie are shown in S5B.

### Supplementary movie S9

3D rendered model (ZEN black software) of Triple knock in A2780 cells (mNeonGreen-Rab11a; mCherry-Rab11b; mTagBFP2-Rab25 DExCon) treated by dox (>94 h). Cells were imaged live while recycling Alexa-647 labelled Transferrin (30 minutes) on glass-bottom dish (MatTek, Ashland, MA, USA; Opti-Klear™ Live Cell Imaging Buffer) coated with 10 μg/ml FN using AiryScan LSM880 (63x). Selected frames from this movie, raw un-rendered, are shown in 4C.

### Supplementary movie S10

3D projection of Triple knock in A2780 cells (mNeonGreen-Rab11a; mCherry-Rab11b; mTagBFP2-Rab25 DExCon) treated by dox (>94 h). Cells were imaged live while recycling Alexa-647 labelled Transferrin (30 minutes) and migrating in <u>C</u>ell <u>D</u>erived <u>M</u>atrix (3D). AiryScan LSM880 (63x); Glass-bottom dish (MatTek, Ashland, MA, USA); Opti-Klear™ Live Cell Imaging Buffer). Selected frames from this movie are shown in 4C.

### Supplementary movie S11

Triple knock in A2780 cells (mNeonGreen-Rab11a; mCherry-Rab11b; mTagBFP2-Rab25 DExCon) treated by dox (>94 h). Cells were imaged live while recycling Alexa-647 labelled Transferrin (30-60 min) on FN-coated 5 mm coverslip. 3D projections and optical sections are shown. Opti-Klear™ Live Cell Imaging Buffer; 3i Lattice LightSheet microscope. Selected frames from this movie are shown in 4F and S5E.

### Supplementary movie S12

Triple knock in A2780 cells (mNeonGreen-Rab11a; mCherry-Rab11b; mTagBFP2-Rab25 DExCon) treated by dox (>94 h) with Alexa-647 labelled Transferrin recycled for 30 min. Cells were imaged fixed on FN-coated 5 mm coverslip. 3D projection animation generated using Imaris Cell Imaging Software. Opti-Klear™ Live Cell Imaging Buffer; 3i Lattice LightSheet microscope. Selected frames from this animation are shown in S5F.

### Supplementary movie S13

Timelapse (Incucyte® S3 system, 20x) of protein expression kinetics of mNeonGreen-Rab11a / mCherry-Rab11b or mNeonGreen-Rab11b / mCherry-Rab11b double DExCon cells (A2780) treated by dox (from time 0). Timelapse covers total 49 h with frame taken every 30 min (approximately 30 min elapsed time per second of the movie); 96 well tissue culture plates (Corning); RPMI fenol-free media. Selected frames from this movie are shown in 5G.

### Supplementary movie S14

Timelapse (Incucyte® S3 system, 20x) of expression and degradation kinetics of miniIAA7-mCherry-Rab11b DExogron cells (A2780, comparison with the classical endogenous mCherry-Rab11b tagging) treated by ±dox ±IAA (from time 0 or cells pre-treated 24h with dox). Timelapse (mCherry channel is shown) covers total 49 h with frame taken every 30 min (approximately 30 min elapsed time per second of the movie); 96 well tissue culture plates (Corning); RPMI fenol-free media. Selected frames from this movie are shown in 6J.

### Supplementary movie S15

Timelapse (Incucyte® S3 system, 20x) of expression and degradation kinetics of miniIAA7-mCherry-Rab11b DExogron / mNeonGreen-Rab11a DExCon cells (A2780, comparison with the classical double endogenous mCherry/mNeonGreen-Rab11b/Rab11a tagging) treated by ±dox ±IAA (from time 0 or cells pre-treated 24h with dox). Timelapse (merge of mCherry/mNeonGreen channel is shown) covers total 49 h with frame taken every 30 min (approximately 30 min elapsed time per second of the movie); 96 well tissue culture plates (Corning); RPMI fenol-free media. Selected frames from this movie are shown in S10A.

### Supplementary movie S16

A2780 (Ctrl; stably expressing AtAFBP2); Rab11s DExCons / DExogrons cells or cells with shRNA anti-Rab11a or b as indicated. Wound healing experiment automatically imaged and analysed in real time by Incucyte® S3 system (blue mask determined based on brightfield image taken at time 0 and every other frame taken after the scratch). Confluent cells were pre-treated for 24 ± dox (250 ng/ml) ± IAA (100 μg/ml), scratch by wound the WoundMaker™ and imaged in RPMI fenol free media ± dox/IAA as indicated. Timelapse (merge of brightfield with highlighted blue mask and true mNeonGreen/mCherry fluorescence) covers total 24 h with frame taken every 1h (approximately 1h elapsed time per second of the movie); ImageLock 96-well Plates; RPMI fenol-free media. Selected frames from this movie are shown in S9A.

### Supplementary movie S17

Timelapse (Incucyte® S3 system, 20x) of expression and degradation kinetics of miniIAA7-mNeonGreen-Rab11a DExogron / mCherry-Rab11b DExCon cells (clone of A2780, comparison with the classical double endogenous mCherry/mNeonGreen-Rab11b/Rab11a tagging) treated by ±dox ±IAA (from time 0 or cells pretreated 24h with dox). Timelapse (merge of mCherry/mNeonGreen channel is shown) covers total 49 h with frame taken every 30 min (approximately 30 min elapsed time per second of the movie); 96 well tissue culture plates (Corning); RPMI fenol-free media. Selected frames from this movie are shown in S11B.

